# A transcription factor quintet orchestrating bundle sheath expression in rice

**DOI:** 10.1101/2024.06.17.599020

**Authors:** Lei Hua, Na Wang, Susan Stanley, Ruth M. Donald, Satish Kumar Eeda, Kumari Billakurthi, Ana Rita Borba, Julian M. Hibberd

**Affiliations:** Department of Plant Sciences, University of Cambridge, Downing Street, Cambridge CB2 3EA, United Kingdom

**Keywords:** *Oryza sativa*, C_4_ rice, Bundle Sheath, Sulfite Reductase, Promoter, cis-elements

## Abstract

C_4_ photosynthesis has evolved in over sixty plant lineages and improves photosynthetic efficiency by ∼50%. One unifying character of C_4_ plants is photosynthetic activation of a compartment such as the bundle sheath, but gene regulatory networks controlling this cell type are poorly understood. In Arabidopsis a bipartite MYC-MYB transcription factor module restricts gene expression to these cells but in grasses the regulatory logic allowing bundle sheath gene expression has not been defined. Using the global staple and C_3_ crop rice we identified the *SULFITE REDUCTASE* promoter as sufficient for strong bundle sheath expression. This promoter encodes an intricate *cis*-regulatory logic with multiple activators and repressors acting combinatorially. Within this landscape we identified a distal enhancer activated by a quintet of transcription factors from the WRKY, G2-like, MYB-related, IDD and bZIP families. This module is necessary and sufficient to pattern gene expression to the rice bundle sheath. Oligomerisation of the enhancer and fusion to core promoters containing Y-patches allowed activity to be increased 220-fold. This enhancer generates bundle sheath-specific expression in *Arabidopsis* indicating deep conservation in function between monocotyledons and dicotyledons. In summary, we identify an ancient, short, and tuneable enhancer patterning expression to the bundle sheath that we anticipate will be useful for engineering this cell type in various crop species.

## Introduction

In plants and animals significant progress has been made in understanding transcription factor networks responsible for the specification of particular cell types. In animals, for example, homeobox transcription factors define the body plan of an embryo (Lewis 1978; Krumlauf 1994), and cardiac cell fate is specified by a collective of five transcription factors comprising Pnr and Doc that act as anchors for dTCF, pMad and Tin (Junion et al. 2012). In plants the INDETERMINATE DOMAIN (IDD) transcription factors work together with SCARECROW and SHORTROOT to specify endodermal formation in the root (Moreno-Risueno et al. 2015; Drapek et al. 2017), PHLOEM EARLY (PEAR) and VASCULAR-RELATED NAC DOMAIN (VND) transcription factors permit production of phloem and xylem vessel respectively (Kubo et al. 2005; Miyashima et al. 2019),and basic helix–loop–helix (bHLH) transcription factors determine differentiation of guard cells (MacAlister et al. 2006; Ohashi-Ito and Bergmann 2006; Pillitteri et al. 2006; Kanaoka et al. 2008). Moreover, transcription factor networks that integrate processes as diverse as responses to external factors such as pathogens and abiotic stresses(Nakashima et al. 2009; Tsuda and Somssich 2015), or internal events associated with the circadian clock (McClung 2006; Nagel and Kay 2012) and hormone signalling (Depuydt and Hardtke 2011; Verma et al. 2016) have also been identified. Transcription factor activity is decoded by short *cis*-acting DNA sequences known as enhancers. The binding of multiple transcription factors to enhancers thus controls transcription and the spatiotemporal patterning of gene expression. For example, the Block C enhancer interacts with the core promoter to activate expression of FLOWERING LOCUS T (FT) in long days (Adrian et al. 2010; Liu et al. 2014), and a distant upstream enhancer controls expression of the *TEOSINTE BRANCHED1* locus in maize responsible for morphological differences compared with the wild ancestor teosinte (Stam et al. 2002; Clark et al. 2006). In contrast to the above examples, transcription factors and cognate *cis*-elements responsible for the operation of cell types in grasses once specified have not been defined (Weber et al. 2016; Schmitz et al. 2022).

Given the increased specialisation of organs evident since the colonisation of land this lack of understanding of gene regulatory networks controlling cell specific gene expression is striking. For example, in the liverwort *Marchantia polymorpha* the photosynthetic thallus contains seven cell types (Wang et al. 2023), while leaves of *Oryza sativa* (rice) and *Arabidopsis thaliana* possess at least fifteen and seventeen populations of cells as defined by single-cell sequencing respectively (Wang et al. 2021). In leaves of these angiosperms, particular cell types are specialised for photosynthesis and so whilst photosynthesis gene expression is induced by light in all major cell types of the rice leaf the response is greater in spongy and palisade mesophyll cells compared with guard, mestome and bundle sheath cells (Swift et al. 2023). In the case of the bundle sheath, these cells carry out photosynthesis, but are specialised to allow water transport from veins to mesophyll, sulphur assimilation and nitrate reduction (Leegood 2008; Aubry et al. 2014b; Hua et al. 2021). And, strikingly in multiple lineages, the bundle sheath has been dramatically repurposed during evolution to become fully photosynthetic and allow the complex C_4_ pathway to operate (Sage 2004).

Compared with the ancestral C_3_ state, plants that use C_4_ photosynthesis operate higher light, water and nitrogen use efficiencies (Makino et al. 2003; Sage 2004; Mitchell and Sheehy 2006). It is estimated that introducing the C_4_ pathway into C_3_ rice would allow a 50% increase in yield (Mitchell and Sheehy 2006; Hibberd et al. 2008), but it requires multiple photosynthesis genes to be expressed in the bundle sheath, including enzymes that decarboxylate C_4_ acids to release CO_2_ around RuBisCO, organic acid transporters, components of the Calvin-Benson-Bassham cycle, RuBisCO activase, and enzymes of starch biosynthesis (Kajala et al. 2011; Aubry et al. 2014a; Ermakova et al. 2021). In summary, although the bundle sheath is found in all angiosperms and associated with multiple processes fundamental to leaf function, the molecular mechanisms responsible for directing expression to this cell type, including in global staple crops, remain undefined. We therefore studied the bundle sheath to better understand the complexity of gene regulatory networks that operate to maintain function of a cell type once it has been specified. Rice was chosen as it a global crop, and identifying how it patterns gene expression to the bundle sheath could facilitate engineering of this cell type.

We hypothesized that analysis of endogenous patterns of gene expression in the rice bundle sheath would allow us to identify a strong and early-acting promoter for this cell type. Once such a promoter was identified we also hypothesised that it could be used to initiate an understanding of the *cis*-regulatory logic that allows gene expression to be patterned to this cell type in grasses. We tested twenty-five promoters from rice genes that transcriptome sequencing indicated were highly expressed in these cells. Of these, four specified preferential expression in the bundle sheath, and one derived from the *SULFITE REDUCTASE* (*SiR*) gene generated strong bundle sheath expression from plastochron 3 leaves onwards. Truncation analysis showed that bundle sheath expression pattern from the *SiR* promoter is mediated by a short distal enhancer and a pyrimidine patch in the core promoter. This bundle sheath module is cryptic until other enhancers acting to both constitutively activate and repress expression in mesophyll cells are removed. The enhancer is composed of a quintet of *cis*-elements recognised by their cognate transcription factors from the WRKY121, GLK2, MYBS1, IDD and bZIP families. These transcription factors act synergistically and are sufficient to drive expression of the strong bundle sheath *SiR* promoter.

## Results

### The *SiR* promoter directs expression to the rice bundle sheath

To identify sequences allowing robust expression in rice bundle sheath cells we used data derived from laser capture microdissection of bundle sheath strands and mesophyll cells from mature leaves. Promoter sequence from seven of the most strongly expressed genes in bundle sheath strands (**Supplemental Figure 1A**) were cloned, fused to the β-glucoronidase (GUS) reporter and transformed into rice. Although five of these promoters (*MYELOBLASTOSIS*, *MYB*; *HOMOLOG OF E. COLI BOLA*, *bolA*; *GLUTAMINE SYNTHETASE 1*, *GS1*; *STRESS RESPONSEIVE PROTEIN*, *SRP*; *ACYL COA BINDING PROTEIN*, *ARP*) led to GUS accumulation, it was restricted to veins **(Supplemental Figure 1B, 1C)**. And, for the *SULFATE TRANSPORTER 3;1* and *3;3* (*SULT3;1* and *SULT3;3*) promoters, no staining was observed (**Supplemental Figure 1B, 1C**). The approach of cloning promoters from bundle sheath strands therefore appeared to be more efficient at identifying sequences capable of driving expression in veins. We therefore optimised a procedure allowing bundle sheath cells to be separated from veins (Hua and Hibberd, 2019) and produced high quality transcriptomes from mesophyll, bundle sheath and vascular bundles (Hua et al. 2021). From these data eighteen genes whose transcripts were more abundant in bundle sheath cells compared with both veins and mesophyll cells were identified (**Supplemental Figure 2A**). When the promoter from each gene was fused to GUS and transformed into rice, those from *ATP-SULFURYLASE 1B*, *ATPS1b*; *SULFITE REDUCTASE*, *SiR*; *HIGH ARSENIC CONTENT1.1*, *HAC1.1*; and *FERREDOXIN*, *Fd* were sufficient to generate expression in the bundle sheath (**Supplemental Figure 2B)**. However, *ATPS1b* and *Fd* also displayed weak activity in the mesophyll, and the *HAC1.1* promoter also led to GUS accumulation in epidermal and vascular cells. Thus, only the *SiR* promoter drove strong expression in the bundle sheath with no GUS detected in other cells (**Supplemental Figure 2B, 2C**). An additional six promoters (*SOLUBLE INORGANIC PYROPHOSPHATASE*, *PPase*; *PLASMA MEMBRANE INTRINSIC PROTEIN1;1*, Os*PIP1;1*; *PLASMA MEMBRANE INTRINSIC PROTEIN1;3*, Os*PIP1;3*; *ACTIN-DEPOLYMERIZING FACTOR*, *ADF*; *PEPTIDE TRANSPORTER PTR2*, *PTR2*; *NITRATE REDUCTASE2*, *NIA2*) generated expression in vascular bundles, and eight promoters produced no staining (**Supplemental Figure 2B, 2C**). In summary, most candidate promoters failed to generate expression that was specific to bundle sheath cells, but the region upstream of the rice *SiR* gene was able to do so. We therefore selected the *SiR* promoter for further characterization.

### The *SiR* promoter drives strong and early expression in bundle sheath cells

Sequence upstream of the *SiR* gene comprising nucleotides −2571 to +42 relative to the predicted translational start site was sufficient to generate expression in the rice bundle sheath. To allow faster analysis of sequences responsible for this output we domesticated the sequence by removing four *Bsa*I and *Bpi*I sites such that it was compatible with the modular Golden Gate cloning system. When this modified sequence was placed upstream of the GUS reporter it also generated bundle sheath preferential accumulation **(Figure 1A)**. Fusion to a nuclear-targeted mTurquoise2 fluorescent protein confirmed that the *SiR* sequence was sufficient to direct expression to bundle sheath cells, and also revealed expression in the longer nuclei of veinal cells **(Figure 1B)**. Expression from the domesticated and non-domesticated sequences was not different **(Figure 1C)**. Compared with 0.58 nmol 4-MU/min/mg protein previously reported for the *Zoysia japonica PHOSPHOENOLCARBOXYKINASE (PCK)* promoter (Emmerling 2018) activity from the *SiR* promoter was at least 36% higher. Designer Transcription Activator-Like Effector (dTALEs) and cognate Synthetic TALE-Activated Promoters (STAPs) amplify expression and allow multiple transgenes to be driven from a single promoter (Brückner et al., 2015; Danila et al., 2022). We therefore tested whether bundle sheath expression mediated by the *SiR* promoter is maintained and strengthened by the dTALE-STAP system. Stable transformants showed bundle sheath specific expression **(Supplemental Figure 3A, 3B)**, and GUS activity was ∼18-fold higher than that from the endogenous *SiR* promoter (**Supplemental Figure 3C**). We conclude that the *SiR* promoter is compatible with the dTALE-STAP system and its activity can be strengthened. We also investigated when promoter activity was first detected during leaf development and discovered that GUS as well as fluorescence from mTurquoise2 were visible in 5-20mm long fourth leaves at plastochron 3 (**Supplemental Figure 4**). This was not the case for the *ZjPCK* promoter even when a dTALE was used to amplify expression (**Supplemental Figure 4**). We conclude that the *SiR* promoter initiates expression in the bundle sheath before the *ZjPCK* promoter, and that it is also able to sustain higher levels of expression in this cell type.

**Figure 1.**
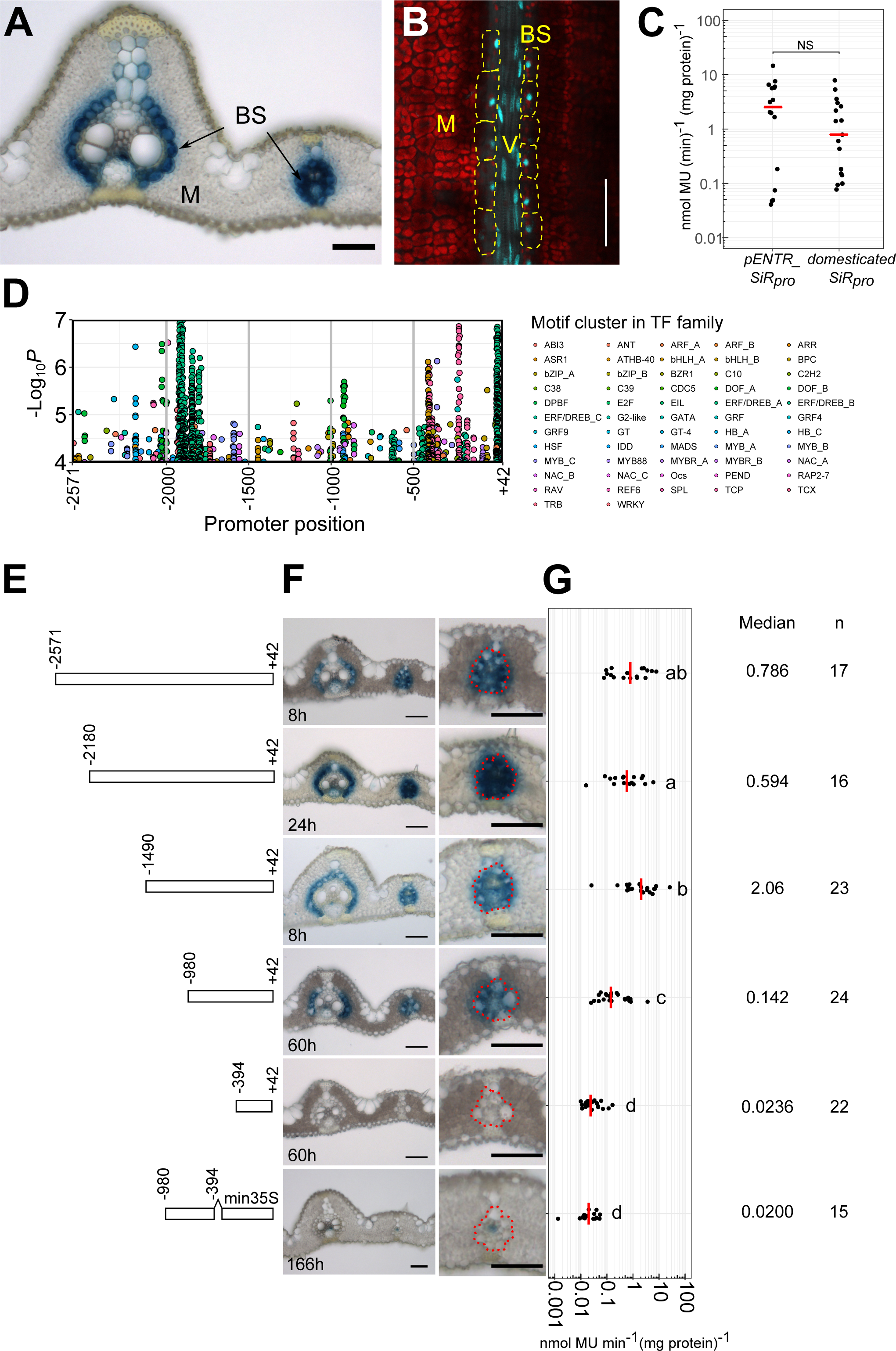
Nucleotides −980 to −394 of the *SiR* promoter are necessary for bundle sheath expression. (A) Domesticated *SiR* promoter generates strong GUS staining in bundle sheath. (B) mTurquoise2 signal driven by the domesticated *SiR* promoter in nuclei (indicated by yellow arrows) of bundle sheath cells (marked by yellow dashed lines) and vein cells in mature leaves, red indicates chlorophyll autofluorescence. **(C)** The fluorometric 4-methylumbelliferyl-β-D-glucuronide (MUG) assay shows no statistically significant difference between the endogenous and domesticated *SiR* promoter activity. **(D)** Landscape of transcription factor binding sites in the *SiR* promoter using the Find Individual Motif Occurrences (FIMO) program. The likelihood of match between 656 plant nonredundant known transcription factor motifs in the *SiR* promoter is shown by transcription factor families (Supplemental table 1). **(E)** Schematics showing 5’ truncations. **(F)** Representative images of leaf cross sections from transgenic lines after GUS staining. Zoomed-in images of lateral veins shown in right panels, the staining duration is displayed in the bottom-left corner, bundle sheath cells highlighted with dashed red line, scale bars = 50 µm. **(G)** Promoter activity determined by the fluorometric 4-methylumbelliferyl-β-D-glucuronide (MUG) assay. Data were subjected to a pairwise Wilcoxon test with Benjamini-Hochberg correction. Lines with differences in activity that were statistically significant (adjusted *P*<0.05) labelled with different letters. Median catalytic rate of GUS indicated with red line, n indicates total number of transgenic lines assessed.

### A distal enhancer and Y-patch necessary for expression in the bundle sheath

The *SiR* promoter contains a highly complex *cis* landscape **(Figure 1D)** comprising at least 638 predicted motifs from 56 transcription factor families (**Supplemental table 1**). We therefore designed a 5’ truncation series to investigate regions necessary for expression in the bundle sheath (**Figure 1E**). Deleting nucleotides −2571 to −2180 and −1490 to −980 led to a statistically significant reduction and then increase in MUG activity respectively but neither truncation abolished preferential accumulation of GUS in the bundle sheath **(Figure 1E-F).** However, when nucleotides −980 to −394 upstream of the predicted translational start site were removed GUS was no longer detectable in bundle sheath cells (**Figure 1E-F**). Consistent with this, MUG assays showed a statistically significantly reduction in activity when these nucleotides were absent **(Figure 1G)**. Thus, nucleotides spanning −980 to −394 of the *SiR* promoter are necessary for bundle sheath specific expression.

To test whether this region is sufficient for bundle sheath specific expression we linked it to the minimal CaMV35S core promoter. Although weak GUS signal was detected in a few veinal cells, this was not the case for the bundle sheath **(Figure 1E-G)**. We conclude that sequence in two regions of the promoter (from −394 to +42 and from −980 to −394) interact to specify expression to the bundle sheath. To better understand this interaction we next generated unbiased 5’ and 3’ deletions. This second deletion series further reinforced the notion that the *SiR* promoter contains a complex *cis*-regulatory landscape. For example, when nucleotides −980 to −829 were removed very weak GUS staining was observed and the MUG assay confirmed that activity was significantly reduced to 11% that of the full-length promoter **(Figure 2, Supplemental Figure 5).** We conclude that nucleotides −980 to −829 from the *SiR* promoter are necessary for tuning expression in the leaf. When nucleotides −829 to −700 were removed GUS appeared in mesophyll cells **(Supplemental Figure 5)**. Truncating nucleotides −613 to −529 abolished GUS accumulation (**Supplemental Figure 5)**. The 3’ deletion that removed nucleotides −251 to +42 also stopped accumulation of GUS in both bundle sheath and mesophyll cells (**Figure 2A-C, Supplemental Figure 5)**. Notably, when the distal region required for bundle sheath expression (−980 to −829) was combined with nucleotides −251 to +42 these two regions were sufficient for patterning to this cell type (**Figure 2**).

**Figure 2.**
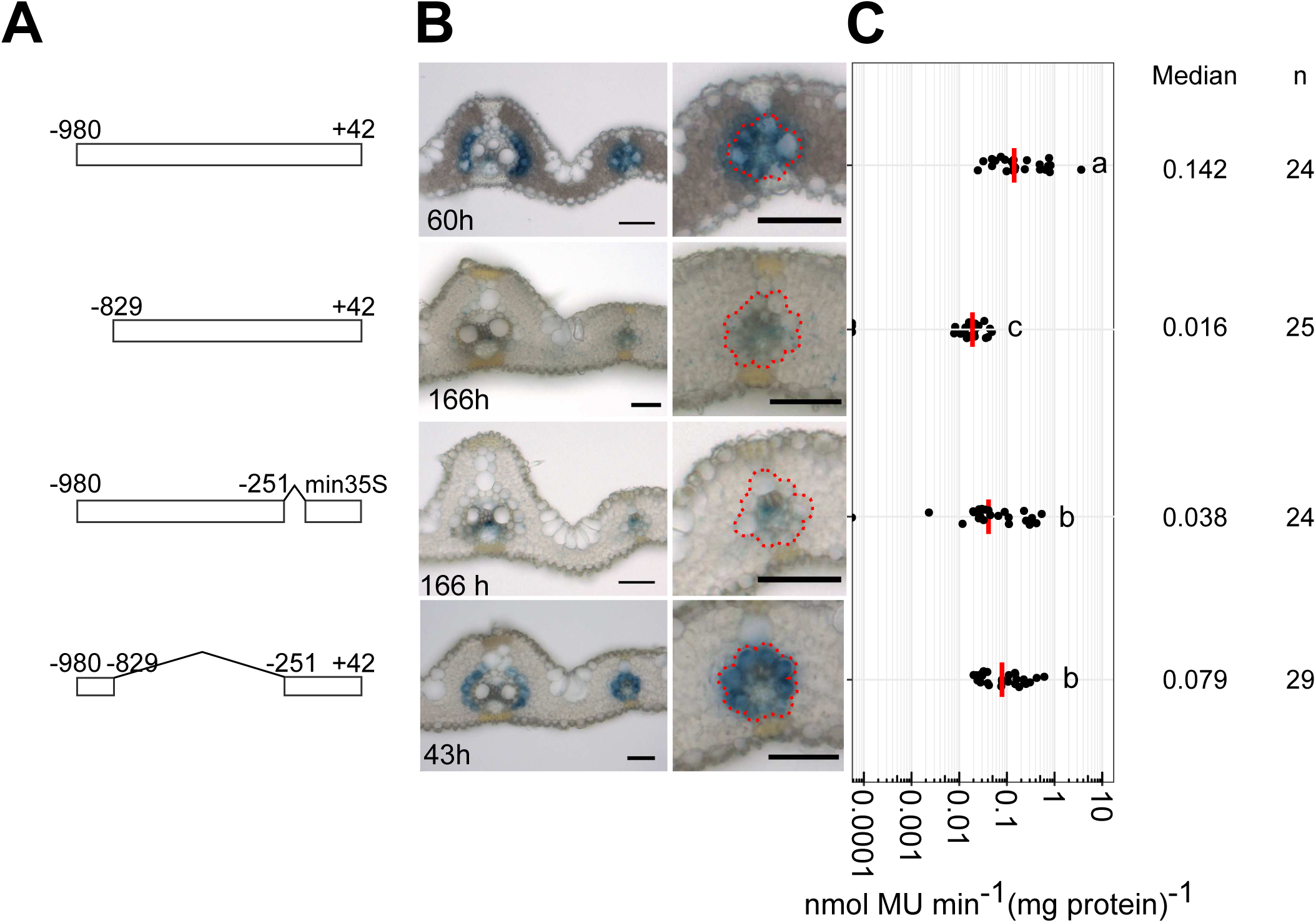
A distal enhancer and the core promoter that are necessary and sufficient for bundle sheath expression. (**A**) Schematics showing deletions of nucleotides −980 to −849 and - 119 to +42. (**B**) Representative image of leaf cross sections of transgenic lines after GUS staining. Zoomed-in images of lateral veins shown in right panels, the staining duration is displayed in the bottom-left corner, bundle sheath cells highlighted with dashed red line, scale bars = 50 µm. (**C**) Promoter activity determined by the fluorometric 4-methylumbelliferyl-β-D-glucuronide (MUG) assay, data subjected to pairwise Wilcoxon test. Lines with differences in activity that were statistically significant (adjusted *P*<0.05) labelled with different letters. Median catalytic rate of GUS indicated with red line, n indicates total number of transgenic lines assessed.

Having identified a region in the *SiR* promoter that was necessary and sufficient for patterning to the bundle sheath, we next used phylogenetic shadowing and yeast one hybrid analysis to better understand the *cis-*elements and *trans*-factors responsible. Analysis of *cis*-elements in the *SiR* promoter that are highly conserved in grasses identified a short region located from nucleotides - 588 to −539 that contained an *ETHYLENE INSENSITIVE3-LIKE 3* (*EIL3*) transcription factor binding site (**Supplemental Figure 6A&B**). Whilst deletion of this motif had no detectable effect of patterning to the bundle sheath (**Supplemental Figure 6C**) the level of expression was reduced (**Supplemental Figure 6D**). We infer that the *EIL3* motif positively regulates activity of the *SiR* promoter but is not responsible for cell specificity. These data are consistent with the promoter truncation analysis that showed nucleotides −613 to −529 containing this motif were not required for bundle sheath specific expression, but instead function as a constitutive activator (**Supplemental Figure 5**). When yeast one hybrid was used to search for transcription factors capable of binding the *SiR* promoter, sixteen were identified (**Supplemental Figure 7A, 7B**). For each, cognate binding sites were present. This included TCP21 and OsOBF1 that can bind to TCP motifs and Ocs/bZIP elements respectively. Consistent with the outcome of deleting the *EIL3* motif, three EIL transcription factors interacted with nucleotides −899 to −500 (**Supplemental Figure 7B, 7C**). Examination of transcript abundance in mature leaves showed that most of these transcription factors were expressed in both bundle sheath and mesophyll cells (**Supplemental Figure 7D**) implying that combinatorial interactions with cell specific factors are likely required for bundle sheath specific expression from the *SiR* promoter.

### The enhancer contains four subregions that simultaneously activate in bundle sheath and repress in mesophyll cells

The truncation analysis above identified two short regions comprising nucleotides −980 to −829 and −251 to +42 that were necessary and sufficient for expression in the rice bundle sheath (**Figure 2, Supplemental Figure 8**). Sequence spanning nucleotides −251 to +42 includes both the annotated 5’ untranslated region but also likely contains core promoter elements (**Supplemental Figure 9A**). Re-analysis of publicly available data identified two major transcription start sites at positions −91 (TSS1) and −41 (TSS2) (**Supplemental Figure 9A**). Although no canonical TATA-box was evident in this region, a TATA-box variant was detected at position −130 (5’-ATTAAA-3’) (Civáň and Švec 2009) that could be responsible for transcription from TSS1. Moreover, upstream of TSS2 is a putative pyrimidine patch (Y-patch) that represents an alternate but common TC-rich core promoter motif in plant genomes (Civáň and Švec 2009) (**Supplemental Figure 9A**). Scanning sequence from −251 to +42 for core promoter elements also identified MTE (Motif Ten Element), BREu (TFIIB Recognition Element upstream) and DCE-S-I (Downstream Core Element S-I) motifs associated with eukaryotic core promoters (**Supplemental Figure 9B**). We therefore assume the region upstream of TSS1 and TSS2 contains the core promoter elements. When consecutive deletions to this sequence were made, statistically significant reductions in MUG activity were evident but there was no impact on accumulation of GUS in the bundle sheath. Interestingly, when the Y-patch was retained but the TATA-box like motif removed, low levels of GUS specific to the bundle sheath were apparent (**Supplemental Figure 9C, 9D, 9E, 9F**), and deletion of the Y-patch completely abolished GUS staining (**Supplemental Figure 9C, 9D, 9E, 9F**). Consistent with the Y-patch being important for bundle sheath expression, when core promoters from other genes (*PIP1;1, NRT1.1A*) containing a Y-patch were linked to the distal enhancer from *SiR* bundle sheath expression was detected **(Figure 3A,3B)**, but this was not the case for genes with only a TATA-box **(Figure 3A,3B)**. GUS activity was higher from the *PIP1;1* core promoter that contains more Y-patches. Overall, we conclude that the TATA-box like motif is not required for expression in the bundle sheath, but the Y-patch is necessary for this patterning and in combination with a distal enhancer comprising nucleotides −980 to −829 it is sufficient for expression in this cell type.

**Figure 3.**
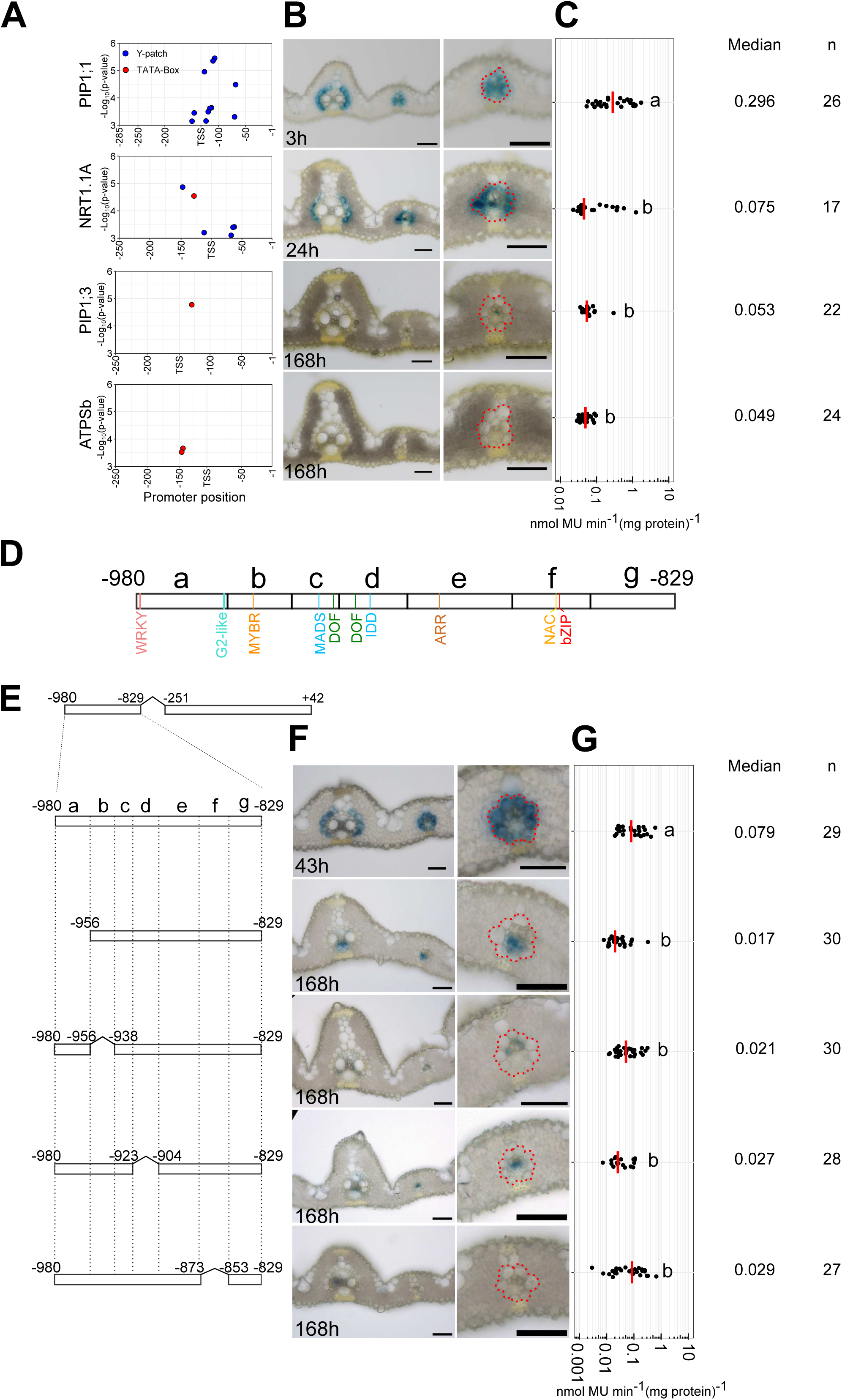
The Y-patch and four distinct regions in the distal enhancer are required for bundle sheath specific expression. **(A-C)** Nucleotides −980 and −829 from the *SiR* promoter pattern expression to the bundle sheath when linked with the *PIP1;1* and *NRT1.1A* core promoters containing Y-patches. **(A)** Prediction of Y-patch and TATA-box sequences in core promoters of *PIP1;1*, *NRT1.1A*, *PIP1;3* and *ATPSb*. **(B)** Representative cross sections of transgenic rice leaves after GUS staining, zoomed-in image of lateral veins shown in the right panel, bundle sheath cells highlighted with white dashed lines, the staining duration is displayed in the bottom-left corner, scale bars = 50 µm. **(C)** Promoter activity determined by the fluorometric 4-methylumbelliferyl-β-D-glucuronide (MUG) assay. **(D)** Schematics showing transcription factor binding sites between nucleotides −980 and −829. **(E)** Schematics showing consecutive deletions between nucleotides - 980 and −829 fused to the GUS reporter. **(F)** Representative images of cross sections from transgenic lines after GUS staining, zoomed-in images of lateral veins shown in right panels, the staining duration is displayed in the bottom-left corner, bundle sheath cells highlighted with red dashed lines, scale bars = 50 µm. **(G)** Promoter activity determined by the fluorometric 4-methylumbelliferyl-β-D-glucuronide (MUG) assay. In **C**&**G**, data were subjected to pairwise Wilcoxon test with Benjamini-Hochberg correction. Lines with differences in activity that were statistically significant (adjusted *P*<0.05) labelled with different letters. Median catalytic rate of GUS indicated with red line, n indicates total number of transgenic lines assessed.

We assessed the distal enhancer for transcription factor binding sites. The FIMO algorithm identified motifs associated with WRKY, G2-like, MYB-related, MADS, DOF, IDD, ARR, SNAC (Stress-responsive NAC) families. PlantPAN (Chow et al. 2018), which includes historically validated *cis-*elements, found an additional Dc3 Promoter Binding Factor (DPBF) binding site for group A bZIP transcription factors **(Figure 3D)**. Seven consecutive deletions spanning this enhancer region and hereafter termed subregions a-g were generated **(Figure 3D)**. Although veinal expression persisted when subregions a, b and d were absent, deletion of subregions a, b, d and f resulted in loss of GUS from bundle sheath cells (**Figure 3E-3G**). MUG analysis showed that deletion of all four regions significantly reduced promoter activity **(Figure 3G)**. In contrast, deletions of nucleotides −938 to −923 (subregion c), −904 to 873 (subregion e), and −853 to −829 (subregion g) had no impact on the patterning **(Supplemental Figure 10)**. The subregions necessary for expression in the bundle sheath contained unique binding sites for WRKY, G2-like, MYB-related, IDD, NAC and bZIP (DPBF) transcription factors. To examine the significance of these regions in the context of full-length *SiR* promoter, consecutive deletions from subregion a to f were generated **(Supplemental Figure 11A)**. Deletion of subregion a, d or f, led to GUS accumulating primarily in mesophyll cells whereas removal of subregion b, c or e, caused GUS staining in both mesophyll cells and bundle sheath cells **(Supplemental Figure 11B)**. No significant changes in GUS activity were observed in these deletion lines **(Supplemental Figure 11C)**. We conclude that that the distal enhancer generates expression in the bundle sheath due to four distinct sub-regions, and that nucleotides between −980 to −853 also function as repressors of mesophyll expression by interacting with nucleotides −829 to −251.

### WRKY, G2-like, MYB-related, IDD and bZIP transcription factors activate the distal enhancer

To gain deeper insight how the distal enhancer operates we employed multiple approaches including transactivation assays, co-expression analysis and site directed mutagenesis. The distal enhancer contained WRKY, G2-like, MYBR, IDD, SNAC and bZIP (DPBF) motifs **(Figure 4A, Supplemental Figure 12)**. We therefore cloned rice transcription factors from each family and used them as effectors in transient assays **(Supplemental Figure 13)**. WRKY121, GLK2, MYBS1, IDD2/3/4/6/10, and bZIP3/4/9/10/11 transcription factors led to the strongest activation of expression from the bundle sheath enhancer **(Figure 4B**, **Supplemental Figure 14A-14D)**, whereas the stress-responsive NAC transcription factors targeting a SNAC motif that overlaps a bZIP (DPBF) motif, activated less strongly than bZIP factors **(Supplemental Figure 14E)**. We therefore conclude that the SNAC motif is not important for activity of the bundle sheath enhancer. Effector assays using pairwise combinations of transcription factors showed synergistic activation from the distal enhancer when GLK2 and IDD3,4,6,10 were co-expressed **(Figure 4C)**.

**Figure 4.**
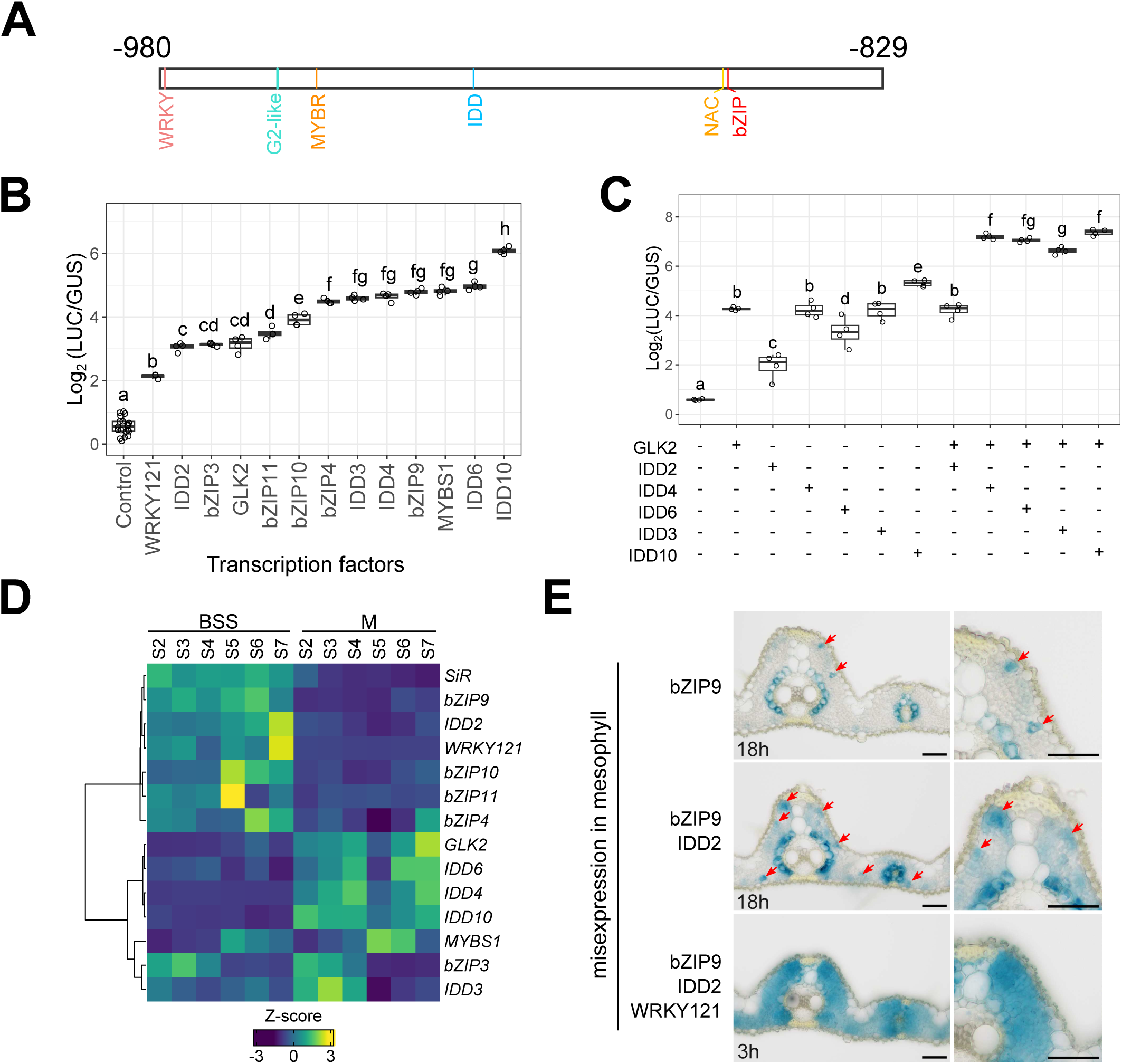
WRKY, G2-like, MYB-related, IDD and bZIP transcription factors interact and activate with the distal enhancer. **(A)** Schematics showing transcription factor binding sites between nucleotides −980 and −829 which are likely required for bundle sheath specific expression. **(B)** Effector assays showing that each transcription factor activates expression from the distal enhancer. **(C)** Effector assays showing synergistic activation from the distal enhancer when GLK2 and IDD3,4,6,10 were co-expressed. Data subjected to pairwise Wilcoxon test with Benjamini-Hochberg correction in B&C. Lines with differences in activity that were statistically significant (adjusted *P*<0.05) labelled with different letters. **(D)** Transcript abundance of transcription factors in bundle sheath strands (BSS) and mesophyll (M) cells during maturation. Leaf developmental stage S2 to S7 represent base of the 4^th^ leaf at the 6th, 8th, 9th, 10th, 13th and 17th day after sowing. (**E**) Representative images of transgenic lines misexpressing WRKY121, IDD2 and bZIP9 in mesophyll cells, staining duration is displayed in the bottom-left corner, zoom-in of mesophyll shown in right panel, red arrows indicate GUS expressing mesophyll cells.

Co-expression analysis using a cell-specific leaf developmental gradient dataset revealed that GLK2, MYBS1 and IDD4,6,10 transcription factors that bind the G2-like, MYB-related and IDD motifs respectively were more abundant in mesophyll cells **(Figure 4D)**. However, the bZIP9, IDD2 and WRKY121 transcription factors strongly correlated with *SiR* transcript abundance and were preferentially expressed in bundle sheath cells **(Figure 4D)**. To test whether bZIP9, IDD2 and WRKY121 are sufficient to pattern *SiR* expression to specific cells, we mis-expressed the single transcription factor bZIP9, both bZIP9 and IDD2, and all three (bZIP9 and IDD2 and WRKY121) in the mesophyll **(Supplemental Figure 15A, 15C, 15E)**. Mis-expression of bZIP alone induced GUS expression from the bundle sheath enhancer in some mesophyll cells **(Figure 4E, Supplemental Figure 15B)**, and mis-expression of both bZIP9 and IDD2 induced greater expression in mesophyll cells **(Figure 4E, Supplemental Figure 15D)**. Strikingly, the expression of bZIP9 and IDD2 and WRKY121 in mesophyll cells fully activated expression in this cell type **(Figure 4E, Supplemental Figure 15F)**. We conclude that one or two transcription factors are weakly sufficient, but all three together effectively interact with the distal enhancer in bundle sheath cells to drive *SiR* expression. We next mutated WRKY, G2-like, MYB-related, IDD, and bZIP motifs. With the exception of the WRKY site that had no statistically robust effect, mutations in each of these motifs diminished or abolished enhancer activity in the bundle sheath **(Figure 5A-5C)**.

**Figure 5.**
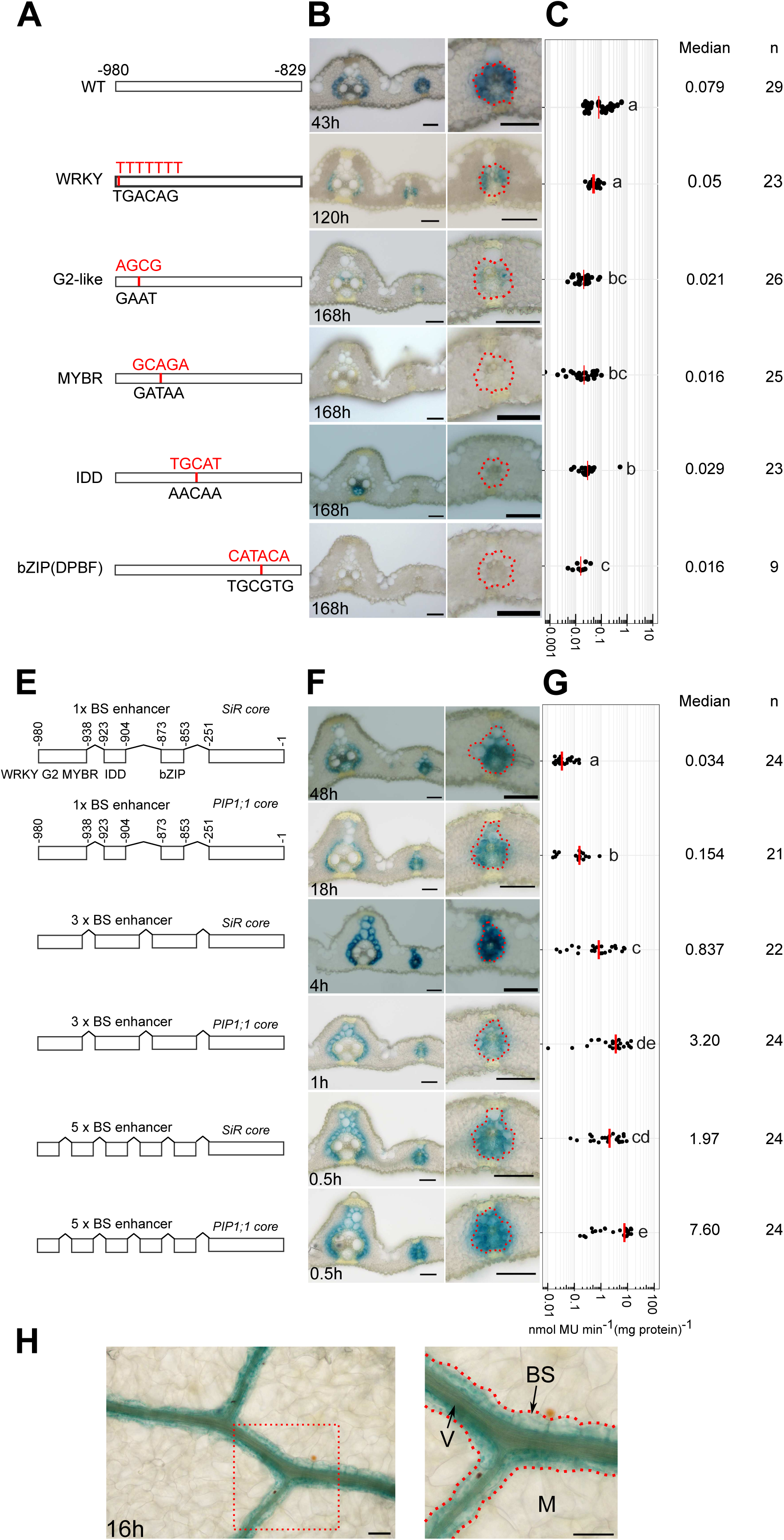
Oligomerisation of bundle sheath enhancer increases bundle sheath expression. Schematics showing site-directed mutagenesis of WRKY, G2-like, MYBR, IDD and bZIP motifs, mutated nucleotides highlighted in red (A), and constructs to test impact of oligomerization of enhancer (E). (B&F) Representative images of cross sections from transgenic lines after GUS staining, zoomed-in images of lateral veins shown in right panel, the staining duration is displayed in the bottom-left corner, bundle sheath cells highlighted with red dashed lines, scale bars = 50 µm. (C&G) Promoter activity determined by the fluorometric 4-methylumbelliferyl-β-D-glucuronide (MUG) assay. Data subjected to pairwise Wilcoxon test with Benjamini-Hochberg correction. Lines with differences in activity that were statistically significant (adjusted *P*<0.05) labelled with different letters. Median catalytic rate of GUS indicated with red line, n indicates total number of transgenic lines assessed, n indicates total number of transgenic lines assessed. (H) Paradermal view of Arabidopsis leaf expressing GUS under the control of 3x BS enhancer combined with OsSiR core promoter, the staining duration is displayed in the bottom-left corner. M indicate mesophyll, BS for bundle sheath, V for vein. Zoomed in images shown on right.

In order to test whether the WRKY, G2-like, MYB-related IDD, and bZIP (DPBF) sites are sufficient to pattern expression to rice bundle sheath cells we concatenated them and fused them to the core promoter of *SiR* (**Figure 5E**). GUS staining was evident in the bundle sheath (**Figure 5F**). Fusion to the *PIP1;1* core promoter maintained bundle sheath expression and resulted in an ∼5 fold increase in activity (**Figure 5E&F)**. Oligomerisation of the enhancer by repeating it three or five times increased bundle sheath specific expression 23 or 58-fold respectively when fused to *SiR* core promoter (**Figure 5E-5G**), and this effect was amplified 90 and 224-fold when fused with the *PIP1;1* core (**Figure 5E-5G**). Synthetic promoters, created by oligomerising this enhancer and combining it with core promoters that contain Y-patches enabled fine-tuning of bundle sheath-specific expression in rice. When an oligomerised version of the enhancer was linked to the *SiR* core promoter and placed in *A. thaliana*, it generated strong expression in bundle sheath cells (**Figure 5H, Supplemental Figure 16**). Collectively our data indicate that transcription factors belonging to the WRKY, G2-like, MYB-related, IDD and bZIP (DPBF) families act cooperatively to decode distinct *cis-*elements in a distal enhancer of the *SiR* promoter, and that this transcription factor collective represent an ancient and highly conserved mechanism allowing bundle sheath specific gene expression in in both monocotyledons and dicotyledons.

## Discussion

### Expression of multiple genes in the rice bundle sheath is not associated with close upstream enhancers

Gene expression is determined by interactions between elements in the core promoter allowing basal levels of transcription (Juven-Gershon and Kadonaga 2009; Haberle and Stark 2018) with more distal *cis*-regulatory modules (Spitz and Furlong 2012; Shlyueva et al. 2014; Ray-Jones and Spivakov 2021). Such *cis*-regulatory modules include enhancers and silencers that act as hubs receiving input from multiple transcription factors and so allow gene expression to respond spatially and temporally to both internal and external stimuli (Li et al. 2007; Buecker and Wysocka 2012). After testing 25 promoters, we discovered that the majority were not capable of driving expression in the rice bundle sheath, and this included ten that generated no detectable activity of GUS in leaves. In all cases we had cloned sequence between −3191 and −960 nucleotides upstream of the predicted translational start site and so these data demonstrate that the core promoter and any enhancers in these regions are not sufficient to direct expression to rice bundle sheath cells. Combined with the paucity of previously reported promoters active in this cell type (Nomura et al. 2005a; Lee et al. 2021) these data argue either for long range upstream enhancers (Studer et al. 2011; Liu et al. 2015; Li et al. 2019; Yan et al. 2019; Zhao et al. 2022) or other regulatory mechanisms being important to specify expression in the bundle sheath. Possibilities include transcription factor binding sites in introns that impact on transcription start site and strongly enhance gene expression (Rose et al. 2008; Gallegos and Rose 2019; Rose 2019), or in exons where because such sequences specify amino acid sequence as well as binding of *trans*-factors, they have been termed duons (Stergachis et al. 2013). Functional analysis showed that duons can pattern expression to the bundle sheath of the C_4_ plant *Gynandropsis gynandra* (Reyna-Llorens et al. 2018), and it is notable that a genome-wide analysis of transcription factor binding sites in grasses revealed genes preferentially expressed in bundle sheath cells tended to contain transcription factor binding sites in their coding sequence (Burgess et al. 2019). It therefore appears possible that gene expression in the bundle sheath is commonly encoded by non-canonical architecture perhaps based on duons rather than more traditional enhancer elements upstream of the core promoter.

Despite the above, we discovered four promoters capable of driving expression in the rice bundle sheath, and each was associated with a gene important in sulphur metabolism. For example, *ATPS*, *SiR* and *Fd* all participate in the first two steps of sulphate reductive assimilation, while *HAC1;1* encodes an arsenate reductase important in the detoxification of arsenate using glutathione that is a product of sulphur assimilation. Collectively, these data further support the notion that the rice bundle sheath cell is specialised in sulphur assimilation (Hua et al. 2021).

### Two distinct genetic networks governing expression in bundle sheath cells

The only other promoter for which both *cis*-elements and *trans*-factors that are necessary and sufficient to pattern bundle sheath expression have been reported is from the dicotyledonous model *A. thaliana*. Here, a bipartite MYC-MYB module upstream of the *MYB76* gene is responsible for this output (Dickinson et al. 2020). MYB76 forms part of a network governing glucosinolate biosynthesis in *A. thaliana*, and so it is notable that the gene regulatory network we report in rice is also associated with sulphur metabolism. However, rather than the bipartite transcription factor network that regulates bundle sheath expression in *A. thaliana,* in rice we report a quintet of transcription factors controlling *SiR* **(Figure 6C&6D)**. The enhancer controlling bundle sheath *SiR* expression in rice comprises four distinct regions recognised by transcription factors belonging to the WRKY, G2-like, MYB-related, IDD and bZIP families **(Figure 6D)**. As loss of the G2-like, MYB-related, IDD and bZIP motifs all reduced expression in the bundle sheath, this implies they act co-operatively - a notion further supported by the fact that GLK2 and IDD3,4,6,10 synergistically activated promoter output in a transient assay. It is of course possible that other motifs in the enhancer such as MADS, DOFs and ARRs act as modulators to tune the level of bundle sheath expression. In fact, single nuclei sequencing of rice and sorghum during photomorphogenesis identified DOFs as important for the evolution of C_4_ gene bundle sheath expression (Swift, Luginbuhl et al. 2023). For the *PIP1;1* and *NRT1.1A* genes, whose transcripts preferentially accumulate in the bundle sheath, the core promoters were not able to generate bundle sheath expression, but they contain a Y-patch and the WRKY, G2-like, MYB-related, IDD and bZIP enhancer is present in intronic sequence **(Supplemental Figure 17).** It is therefore possible that this regulatory system controls their expression. Moreover, for two promoters from rice (*Fd* and *HAC1;1*) and two from other species (*ZjPCK* and *FtGLDP*) that are sufficient to drive expression to the bundle sheath contain Y-patches and the cognate *cis*-elements for WRKY, G2-like, MYB-related, IDD and bZIP transcription factors (**Supplementary Figure 17**).

**Figure 6.**
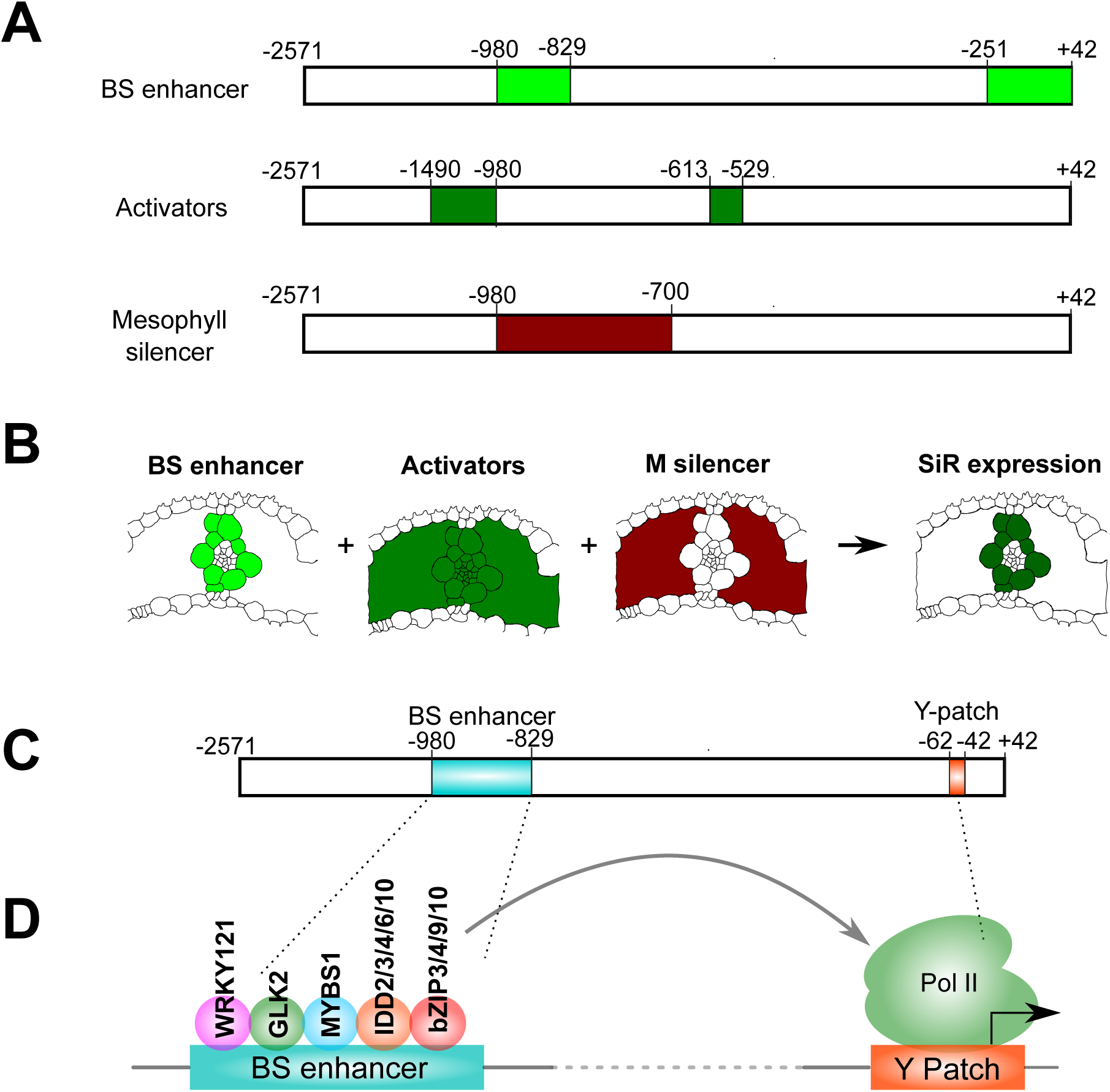
Model of mechanism underpinning bundle sheath expression from *SiR* promoter. **(A)** Schematic with location of Bundle Sheath (BS) enhancer, constitutive activator and mesophyll repressor. (B) Bundle sheath expression is a result of the enhancer, constitutive activators and mesophyll repressor acting in concert. Schematic indicating how the enhancer operates within a broader *cis*-regulatory landscape. (C&D) Model depicting transcription factors and cognate *cis*-elements responsible for bundle sheath expression.

The distal enhancer in the *SiR* promoter operates in conjunction with the core promoter that contains two transcription start sites, one with an upstream TATA-box and the other a TC-rich element known as a pyrimidine (Y) patch **(Supplemental Figure 9A)**. The TATA-box is found in metazoans and plants and allows recognition by the pre-initiation complex (Smale and Kadonaga 2003), but in plants computational analysis showed that many promoters lack a TATA-box and instead contain a Y-patch (Yamamoto et al. 2007a, 2007b; Bernard et al. 2010). These genes tend to be relatively steadily expressed and associated with protein metabolism (Bernard et al., 2010), and presence of a Y-patch can increase core promoter strength (Jores et al. 2021). For *SiR,* whilst the TATA-box is not required, the Y-patch is needed for expression in the bundle sheath. Notably, core promoters with a higher number or longer Y-patches tended to drive stronger expression, and showed that in plants cell specific gene expression can be tuned by selecting different core promoters.

The regulatory network comprising the Y-patch and distal enhancer enabling bundle sheath expression of *SiR* is embedded within a complex *cis*-regulatory landscape with distinct regions encoding activating and repressing activities (**Figure 6A&6B**). For example, the distal enhancer (nucleotides −980 to −829) activating expression in the bundle sheath overlaps with sequence (nucleotides −900 to −700) that suppresses expression in the mesophyll. Notably, the distal enhancer is both essential for mesophyll repression and also sufficient to drive bundle sheath specific expression (**Figure 6A**). In addition to controlling cell specificity, this complexity likely also facilitates the tuning of expression to environmental conditions. For instance, the *EIL* motif (position −572 to −552) is recognised by *ETHYLENE-INSENSITIVE LIKE* transcription factors that respond to sulphur deficiency (Maruyama-Nakashita et al. 2006; Dietzen et al. 2020). As transcripts encoding *EIL* accumulate in both bundle sheath and mesophyll cells in response to sulphate deficiency it seems likely that transcription factors repressing expression in the mesophyll respond in a dynamic manner. In addition to *EIL*, the yeast one hybrid analysis identified seven other families of transcription factor families that can bind the *SiR* promoter. Many play documented roles during abiotic or biotic stress, with for example OBF1, ERF3, NAP and FLP acting during low-temperature or drought responses (Shimizu et al. 2005; Zhang et al. 2013, 2022; Chen et al. 2014; Qu et al. 2022), while TCP21, EREBP1, ERF3, ERF72, ERF83 are involved in both abiotic and biotic stress (Lin et al. 2007; Jisha et al. 2015; Zhang et al. 2016, 2018; Tezuka et al. 2019; Jung et al. 2021). Consistent with previous *in silico* analysis (Kurt et al. 2022) the presence of multiple AP2/ERF and EIL transcription factors binding sites suggests that *SiR* is likely subject to control from ethylene signalling (Binder 2020) and also of transcription factors that respond to ABA and jasmonic acid biosynthesis and signalling (Yaish et al. 2010; Chen et al. 2014; Jisha et al. 2015; Zhang et al. 2016, 2018). Together this implies that multiple phytohormone signalling pathways converge on the *SiR* promoter. These data are similar to those reported for the *SHORTROOT* promoter in *A. thaliana* roots where a complex network of activating and repressing *trans*-factors also tunes expression (Sparks et al. 2016). It is also notable that the architecture we report for the bundle sheath enhancer of *SiR* appears of similar complexity to the collective of five transcription factors used to specify cardiac mesoderm in *Drosophila melanogaster* and vertebrates (Junion et al., 2012). For the five transcription factors that bind the cardiac mesoderm enhancer, the order and positioning of motifs (motif grammar) is flexible. However, this is not always the case, with for example output from the human *interferon-beta* (*INF-*β) enhancer demanding a conserved grammar (Thanos and Maniatis 1995; Panne 2008). Further work will be needed to determine if the bundle sheath enhancer reported here for rice is more similar to one of these models, or indeed, as reported for the Drosophila *eve stripe 2* enhancer, operates as a billboard in different tissues to determine patterning of expression (Kulkarni and Arnosti 2003).

### Using the *SiR* promoter to engineer the rice bundle sheath

In addition to bundle sheath cells being important for sulphur assimilation (Leegood 2008; Aubry et al. 2014b; Hua et al. 2021) they have also been implicated in nitrate assimilation, the control of leaf hydraulic conductance and solute transport (Hua et al. 2021) and the systemic response to high light (Xiong et al. 2021). Moreover, in one of the most striking examples of a cell type being repurposed for a new function, bundle sheath cells have repeatedly been rewired to allow the evolution of C_4_ photosynthesis (Sage 2004). To engineer these diverse processes, specific and tuneable promoters for this cell are required. However, identification of sequence capable of driving specific expression to bundle sheath strands has previously been limited to *A. thaliana* and C_4_ species. For example, the *SCARECROW* (Cui et al. 2014), *SCL23* (Cui et al. 2014), *SULT2;2* (Kirschner et al. 2018) and *MYB76* promoters (Dickinson et al. 2020) are derived from *A. thaliana*, whilst the *Glycine Decarboxylase P-protein (GLDP*) promoter is from the C_4_ dicotyledon *Flaveria trinervia* (Engelmann et al. 2008; Wiludda et al. 2012). In rice, only the C_4_ *Zoysia japonica PCK* and the C_4_ *Flaveria trinervia GLDP* promoters are known to pattern expression to the bundle sheath (Nomura et al. 2005c; Lee et al. 2021). Both are capable of conditioning expression in this cell type, but are weak, turn on late during leaf development and the molecular basis underpinning their ability to restrict expression to the bundle sheath has not been defined. It has therefore not been possible to rationally design or tune expression to this important cell type in rice. The architecture of the *SiR* promoter we report here now provides an opportunity to engineer the bundle sheath.

In summary, from analysis of the ∼2600 nucleotide *SiR* promoter we identify an enhancer comprising 81 nucleotides that with the Y-patch is sufficient to drive expression to bundle sheath cells. Moreover, we show that output can be tuned via two approaches. First, oligomerising the distal enhancer can drastically increase expression. Second, combining it with different core promoters achieved the same output, and correlated with length of the Y-patch present. Our identification of a minimal promoter that drives expression in bundle sheath cells of rice now provides a tool to allow this important cell type to be manipulated. Cell specific manipulation of gene expression has many perceived advantages. For example, when constitutive promoters have been used to drive gene expression gene silencing and reduction of plant fitness due to metabolic penalties (Glick 1995; Que et al. 1997). In contrast, tissue specific promoters allow targeted gene expression either spatially or at particular developmental stages and so allow increased precision in trait engineering (Kummari et al. 2020). The *SiR* promoter and the bundle sheath *cis*-regulatory module that we identify thus provide insights into mechanisms governing cell specific expression in rice, and may also contribute to our ability to engineer and improve cereal crops.

## Materials and methods

### Plant material and growth conditions

Kitaake (*O. sativa* ssp. *japonica*) was transformed using *Agrobacterium tumefaciens* with the following modifications as described previously (Hiei et al., 2008). Mature seeds were sterilized with 2.5% (v/v) sodium hypochlorite for 15 mins, and calli were induced on NB medium with 2 mg/L 2,4-D at 30 °C in darkness for 3-4 weeks. Actively growing calli were then co-incubated with *A. tumefaciens* strain LBA4404 in darkness at 25°C for 3 days, they were selected on NB medium supplied with 35 mg/L hygromycin B for 4 weeks, and those that proliferated placed on NB medium with 10 mg/L hygromycin B for 4 weeks at 28 °C under continuous light. Plants resistant to hygromycin were planted in 1:1 mixture of topsoil and sand and placed in a greenhouse at the Botanic Gardens, University of Cambridge under natural light conditions but supplemented with a minimum light intensity of 390 μmol m^−2^ s^−1^, a humidity of 60%, temperatures of 28°C and 23°C during the day and night respectively, and a photoperiod of 12 h light, 12 h dark. Subsequent generations were grown in a growth cabinet in 12 h light/12 h dark, at 28 °C, a relative humidity of 65%, and a photon flux density of 400 μmol m^−2^ s^−1^.

### Cloning and construct preparation, and motif analysis

The 2613-bp promoter DNA fragment of *SULFITE REDUCTASE* (*SiR,* MSU7 ID: LOC_Os05g42350, RAP-DB ID: Os05g0503300) was originally amplified from Kitaake genomic DNA, with forward primer (5’-3’) “CACCATGCTTGACCATGTGGACTC” and reverse primer (5’-3’) “ACGGAACCCGTGGAACTC”. Gel-purified PCR product was cloned into a Gateway pENTR^TM^ vector to generate *pENTR-SiRpro* using pENTR™/D-TOPO™ Cloning Kit (Invitrogen), then the promoter was recombined into the pGWB3 expression vector and fused with *GUS* gene using LR reaction. The resultant vector was transformed into *A. tumefaciens* strain LBA4404 and transformed into Kitaake. To engineer the *SiRpro* such that it is compatible with the Golden gate system, four *Bsa*I or *Bpi*I restriction enzyme recognition sites at −214, −298, −1468, and −2309 were mutated from T to A and cloned into the *pAGM9121* vector using Golden Gate level 0 cloning reactions and then into a level 0 PU module EC14328, which was used for driving *kzGUS* (intronless GUS) and *H2B-mTurquoise2* reporter genes via Golden gate reaction and using *Tnos* as a terminator. A five prime deletion series was generated using EC14328 as the template and prepared as level 0 PU modules, and a three prime deletion series prepared as level 0 P modules. The minimal *CaMV35S* promoter was used as the U module, and they were linked with *kzGUS* and terminated with *Tnos*.

To test *SiRpro* in the dTALE/STAP system, the 42-bp coding region were excluded and the 2571-bp resultant fragment placed into a level 0 PU module EC14330 and was used to drive *dTALE1*. Two reporters were used. For the GUS reporter *kzGUS* was linked with *STAP62* and terminated with *Tnos*. In the fluorescent reporter construct, a chloroplast targeting peptide fused to *mTurquoise 2* was linked with *STAP 4* and terminated with *Tact2*. In both constructs, *pOsAct1* driving *HYG* (Hygromycin resistant gene) was terminated with *Tnos* and used as the selection marker during rice transformation.

The Find Individual Motif Occurrences (FIMO) tool (Grant et al. 2011) from the Multiple Em for Motif Elucidation (MEME) suite v.5.4.0 (Bailey et al. 2015) was used to search for individual motifs within the promoter sequences using default parameters with “--thresh” of “1e-3”. Position weight matrix of 656 non-redundant plant motifs and 13 RNA polymerase II (POLII) core promoter motifs were obtained from JASPAR (https://jaspar.elixir.no/downloads/) (Fornes et al. 2020). To cluster the transcription factor binding motifs, the RSAT matrix-clustering tool (Castro-Mondragon et al. 2017) was run on all 656 non-redundant plant motifs using the default parameters, which yield 51 motif clusters, these clusters were further divided based on transcription factor families (Supplemental table 2).

### Analysis of GUS and fluorescent reporters

In all cases, to account for position effects associated with transformation via *A. tumefaciens*, multiple T_0_ lines were assessed for each construct. GUS staining was performed as described previously (Jefferson et al., 1987) with the following minor modifications. Leaf tissue was fixed in 90% (v/v) acetone overnight at 4 °C after washing with 100 mM phosphate buffer (pH 7.0). Leaf samples were transferred into 1 mg/ml 5-bromo-4-chloro-3-indolyl glucuronide (X-Gluc) GUS staining solution, subjected to 2 mins vacuum infiltration 5 times, and then incubated at 37 °C for between 1 and 168 hours. Chlorophyll was cleared further using 90% (v/v) ethanol overnight at room temperature. Cross sections were prepared manually using a razor blade and images were taken using an Olympus BX41 light microscopy. Quantification of GUS activity was performed using a fluorometric MUG assay (Jefferson et al., 1987). ∼200 mg mature leaves from transgenic plants were frozen in liquid nitrogen and ground into fine powder with a Tissuelyser (Qiagen). Soluble protein was extracted in 1 mL of 50 mM phosphate buffer (pH 7.0) supplemented with 0.1% [v/v] Triton X-100 and cOmplete™ Protease Inhibitor Cocktail (half tablet per 50 mL). Protein concentration then determined using a Qubit protein assay kit (Invitrogen). The MUG fluorescent assay was performed in duplicates with 20 µl protein extract in MUG assay buffer (50mM phosphate buffer (pH 7.0), 10 mM EDTA-Na_2_, 0.1% [v/v] Triton X-100, 0.1% [w/v] N-lauroylsarcosine sodium, 10 mM DTT, 2 mM 4-methylumbelliferyl-β-D-glucuronide (MUG)) in a 200 µl total volume. The reaction was conducted at 37 ^◦^C in GREINER 96 F-BOTTOM microtiter plate using a CLARIOstar plate reader. 4-Methylumbelliferone (4-MU) fluorescence was recorded every 2 minutes for 20 cycles with excitation at 360 nm and emission detected at 450 nm. 4-MU concentration was determined based on a standard curve of ten 4-MU standards placed in the same plate. GUS enzymatic rates were calculated by averaging the slope of MU production from each of the duplicate reactions.

In order to visualize mTurquoise2 mature leaves were dissected into 2-cm sections, leaf epidermal cells were removed by scraping the leaf surface with a razor blade and then mounted with deionized water. 5-mm and middle sections of 2-cm young tissue of the fourth leaves were dissected and mounted with deionized water directly. Imaging was then performed using a Leica TCS SP8 confocal laser-scanning microscope using a 20x air objective. mTurquoise2 fluorescence was excited at 442 nm with emission at 471–481 nm, chlorophyll autofluorescence was excited at 488 nm with emission at 672–692 nm.

### Yeast one hybrid, protoplast isolation and transactivation assay

The yeast one hybridisation assay was performed by Hybrigenics (https://www.hybrigenics-services.com/). Fragments were synthesized and used as bait. Rice leaf and root cDNA libraries were used as prey. The number of clones screened and concentration of 3-AT were as follows: fragment 1, 70.2 million clones screened with 0 mM 3-AT; fragment 2, 61.5 million clones screened with 0 mM 3-AT; fragment 3, 68.4 million clones screened with 20 mM 3-AT; fragment 4, 57.4 million clones screened with 100 mM 3-AT; fragment 5, 94.2 million clones screened with 200 mM 3-AT.

Rice leaf protoplasts and PEG-mediated transformation were performed as described previously (Page et al. 2019). Golden gate level 1 modules for transformation were isolated using ZymoPURE™ II Plasmid Midiprep Kit, *ZmUBIpro::GUS-Tnos* was used as transformation control. Transcription factor coding sequences were amplified using rice leaf cDNA, with *Bsa*I and *Bpi*I sites mutated, and cloned into Golden gate SC level 0 modules. They were assembled into a level 1 module with a *ZmUBIpro* promoter and *Tnos* terminator module. Nucleotides −980 to −829 with the endogenous core promoter (nucleotide −250 to +42) were fused with the *LUC* reporter to generate output of transcription activity. In each transformation, 2 µg of transformation control plasmids, 5 µg of reporter plasmids, and 5 µg of effector plasmids per transcription factor were combined and mixed with 170 µl protoplasts. After incubation on the benchtop for overnight protein was extracted using passive lysis buffer, GUS activity was determined with 20 μl of protein sample and MUG fluorescent assay as described above, LUC activity was measured with 20 μl of protein sample and 100 μl of LUC assay reagent (Promega) using Clariostar plate reader. Transcriptional activity from the promoter was calculated as LUC luminescence/rate of MUG accumulation.

## Acknowledgements

This work was funded by the Bill and Melinda Gates Foundation C_4_ Rice grant awarded to the University of Oxford (2015-2019 - OPP1129902, and 2019-2024 INV-002970. For the purpose of open access, the authors have applied a Creative Commons Attribution (CC BY) licence to any Author Accepted Manuscript version arising from this submission.

## Author contributions

L.H. and J.M.H. conceived the work. J.M.H. guided execution of experiments and oversaw the project. L.H., N.W., S.S., R.D., K.B., and A.R.B. did the experiments and analysed the data. L.H. and J.M.H. wrote the manuscript with input from all authors.

## Declaration of interests

The authors declare no competing interests.

## Supplementary Figure legend titles

**Supplemental Figure 1.** Analysis of seven rice promoters identified after analysis of transcripts that accumulate preferentially in bundle sheath strands.

**Supplemental Figure 2.** Identification of bundle sheath specific promoters after analysis of transcripts that accumulate preferentially in Kitaake bundle sheath compared with mesophyll cells.

**Supplemental Figure 3.** The domesticated *SiR* promoter combined with the dTALE/STAP system drives strong expression in the rice bundle sheath.

**Supplemental Figure 4.** The rice *SiR* promoter drives expression in bundle sheath cells earlier than the *Zoysia japonica PCK* promoter.

**Supplemental Figure 5.** Impact of 5’ and 3’ deletions between on patterning of GUS from the *SiR* promoter.

**Supplemental Figure 6.** The evolutionally conserved *ETHYLENE INSENSITIVE3-LIKE* (*EIL*) binding site regulates the expression level but not cell specificity of the *SiR* promoter.

**Supplemental Figure 7.** Identification of transcription factors interacting with the *SiR* promoter.

**Supplemental Figure 8. N**ucleotides −980 to −829 in combination with nucleotide −251 to +42 produce bundle sheath specific expression.

**Supplemental Figure 9.** Nucleotides −251 to −1 likely serve as a core promoter.

**Supplemental Figure 10.** Subregions in the distal enhancer not required for bundle sheath specific expression between nucleotides −980 to −829.

**Supplemental Figure 11.** Regions between nucleotides −980 and −829 that in combination with nucleotides −828 to −252 repress mesophyll expression.

**Supplemental Figure 12.** Nucleotide sequence in the distal enhancer.

**Supplemental Figure 13.** Transcription factors used in transactivation assay.

**Supplemental Figure 14.** Effector assay showing the effect of WRKY (A), G2-like (B), IDD (C), MYB-related (D), SNAC and bZIP (E) transcription factors on the distal enhancer.

**Supplemental Figure 15.** Impact of mis-expression of bZIP9, IDD2 and WRKY121 in mesophyll cells on GUS expression pattern driven by bundle sheath enhancer.

**Supplemental Figure 16.** The bundle sheath enhancer produces bundle sheath specific expression in Arabidopsis leaves.

**Supplemental Figure 17.** WRKY, G2-like, MYB-related, IDD and bZIP transcription factor binding sites identified by FIMO program in *Fd*, *HAC1;1*, *PIP1.1*, *NRT1.1A*, *ZjPCK* and *FtGLDP* promoters.

## Notes

### Competing Interest Statement

The authors have declared no competing interest.

### Summary of Updates

Some labels in Figures disappeared in conversion

## References

Adrian J, Farrona S, Reimer JJ, Albani MC, Coupland G, and Turck F. Cis-regulatory elements and chromatin state coordinately control temporal and spatial expression of FLOWERING LOCUS T in arabidopsis. Plant Cell. 2010:22(5):1425–1440. 10.1105/tpc.110.074682

Aubry S, Kelly S, Kümpers BMCC, Smith-Unna RD, and Hibberd JM. Deep Evolutionary Comparison of Gene Expression Identifies Parallel Recruitment of Trans-Factors in Two Independent Origins of C_4_ Photosynthesis. PLoS Genet. 2014a:10(6):e1004365. 10.1371/journal.pgen.1004365

Aubry S, Smith-Unna RD, Boursnell CM, Kopriva S, and Hibberd JM. Transcript residency on ribosomes reveals a key role for the Arabidopsis thaliana bundle sheath in sulfur and glucosinolate metabolism. Plant Journal. 2014b:78(4):659–673. 10.1111/tpj.12502

Bailey TL, Johnson J, Grant CE, and Noble WS. The MEME Suite. Nucleic Acids Res. 2015:43:39–49. 10.1093/nar/gkv416

Bernard V, Brunaud V, and Lecharny A. TC-motifs at the TATA-box expected position in plant genes: A novel class of motifs involved in the transcription regulation. BMC Genomics. 2010:11(1):1–15. 10.1186/1471-2164-11-166/TABLES/4

Binder BM. Ethylene signaling in plants. J Biol Chem. 2020:295(22):7710. 10.1074/JBC.REV120.010854

Buecker C and Wysocka J. Enhancers as information integration hubs in development: lessons from genomics. Trends Genet. 2012:28(6):276. 10.1016/J.TIG.2012.02.008

Burgess SJ, Reyna-Llorens I, Stevenson SR, Singh P, Jaeger K, and Hibberd JM. Genome-Wide Transcription Factor Binding in Leaves from C3 and C4 Grasses. Plant Cell. 2019:31(10):2297–2314. 10.1105/TPC.19.00078

Castro-Mondragon JA, Jaeger S, Thieffry D, Thomas-Chollier M, and Van Helden J. RSAT matrix-clustering: dynamic exploration and redundancy reduction of transcription factor binding motif collections. Nucleic Acids Res. 2017:45(13):e119–e119. 10.1093/NAR/GKX314

Chen X, Wang Y, Lv B, Li J, Luo L, Lu S, Zhang X, Ma H, and Ming F. The NAC family transcription factor OsNAP confers abiotic stress response through the ABA pathway. Plant Cell Physiol. 2014:55(3):604– 619. 10.1093/PCP/PCT204

Chow C-N, Lee T-Y, Hung Y-C, Li G-Z, Tseng K-C, Liu Y-H, Kuo P-L, Zheng H-Q, and Chang W-C. PlantPAN3.0: a new and updated resource for reconstructing transcriptional regulatory networks from ChIP-seq experiments in plants. Nucleic Acids Res. 2018. 10.1093/nar/gky1081

Civáň P and Švec M. Genome-wide analysis of rice (Oryza sativa L. subsp. japonica) TATA box and Y Patch promoter elements. Genome. 2009:52(3):294–297. 10.1139/G09-001/SUPPL_FILE/G09-001.DOC

Clark RM, Wagler TN, Quijada P, and Doebley J. A distant upstream enhancer at the maize domestication gene tb1 has pleiotropic effects on plant and inflorescent architecture. Nat Genet. 2006:38(5):594–597. 10.1038/ng1784

Cui H, Kong D, Liu X, and Hao Y. SCARECROW, SCR-LIKE 23 and SHORT-ROOT control bundle sheath cell fate and function in Arabidopsis thaliana. The Plant Journal. 2014:78(2):319–327. 10.1111/TPJ.12470

Danila F, Schreiber T, Ermakova M, Hua L, Vlad D, Lo SF, Chen YS, Lambret-Frotte J, Hermanns AS, Athmer B, et al. A single promoter-TALE system for tissue-specific and tuneable expression of multiple genes in rice. Plant Biotechnol J. 2022. 10.1111/PBI.13864

Depuydt S and Hardtke CS. Hormone signalling crosstalk in plant growth regulation. Curr Biol. 2011:21(9). 10.1016/J.CUB.2011.03.013

Dickinson PJ, Kneřová J, Szecówka M, Stevenson SSR, Burgess SJ, Mulvey H, Bågman A-MM, Gaudinier A, Brady SM, and Hibberd JM. A bipartite transcription factor module controlling expression in the bundle sheath of Arabidopsis thaliana. Nat Plants. 2020:6(12):1468–1479. 10.1038/s41477-020-00805-w

Dietzen C, Koprivova A, Whitcomb SJ, Langen G, Jobe TO, Hoefgen R, and Kopriva S. The Transcription Factor EIL1 Participates in the Regulation of Sulfur-Deficiency Response. Plant Physiol. 2020:184(4):2120–2136. 10.1104/pp.20.01192

Drapek C, Sparks EE, and Benfey PN. Uncovering Gene Regulatory Networks Controlling Plant Cell Differentiation. Trends in Genetics. 2017:33(8):529–539. 10.1016/J.TIG.2017.05.002

Emmerling J. Studies into the Regulation of C4 Photosynthesis – Towards Factors Controlling Bundle Sheath Expression and Kranz Anatomy Development. Doctoral thesis, Heinrich Heine University, Düsseldorf. 2018:(September).

Engelmann S, Wiludda C, Burscheidt J, Gowik U, Schlue U, Koczor M, Streubel M, Cossu R, Bauwe H, and Westhoff P. The Gene for the P-Subunit of Glycine Decarboxylase from the C4 Species Flaveria trinervia: Analysis of Transcriptional Control in Transgenic Flaveria bidentis (C4) and Arabidopsis (C3). Plant Physiol. 2008:146(4):1773–1785. 10.1104/PP.107.114462

Ermakova M, Arrivault S, Giuliani R, Danila F, Alonso-Cantabrana H, Vlad D, Ishihara H, Feil R, Guenther M, Borghi GL, et al. Installation of C _4_ photosynthetic pathway enzymes in rice using a single construct. Plant Biotechnol J. 2021:19(3):575–588. 10.1111/pbi.13487

Fornes O, Castro-Mondragon JA, Khan A, Van Der Lee R, Zhang X, Richmond PA, Modi BP, Correard S, Gheorghe M, Baranašić D, et al. JASPAR 2020: Update of the open-Access database of transcription factor binding profiles. Nucleic Acids Res. 2020:48(D1):D87–D92. 10.1093/nar/gkz1001

Gallegos JE and Rose AB. An intron-derived motif strongly increases gene expression from transcribed sequences through a splicing independent mechanism in Arabidopsis thaliana. Scientific Reports 2019 9:1. 2019:9(1):1–9. 10.1038/s41598-019-50389-5

Glick BR. Metabolic load and heterologous gene expression. Biotechnol Adv. 1995:13(2):247–261. 10.1016/0734-9750(95)00004-A

Grant CE, Bailey TL, and Noble WS. FIMO: Scanning for occurrences of a given motif. Bioinformatics. 2011:27(7):1017–1018. 10.1093/bioinformatics/btr064

Haberle V and Stark A. Eukaryotic core promoters and the functional basis of transcription initiation. Nat Rev Mol Cell Biol. 2018:19(10):621. 10.1038/S41580-018-0028-8

Hibberd JM and Covshoff S. The Regulation of Gene Expression Required for C _4_ Photosynthesis. Annu Rev Plant Biol. 2010:61(1):181–207. 10.1146/annurev-arplant-042809-112238

Hibberd JM, Sheehy JE, and Langdale JA. Using C4 photosynthesis to increase the yield of rice-rationale and feasibility. Curr Opin Plant Biol. 2008:11(2):228–231. 10.1016/j.pbi.2007.11.002

Hua L, Stevenson SR, Reyna-Llorens I, Xiong H, Kopriva S, and Hibberd JM. The bundle sheath of rice is conditioned to play an active role in water transport as well as sulfur assimilation and jasmonic acid synthesis. The Plant Journal. 2021:107(1):268–286. 10.1111/TPJ.15292

Jisha V, Dampanaboina L, Vadassery J, Mithöfer A, Kappara S, and Ramanan R. Overexpression of an AP2/ERF Type Transcription Factor OsEREBP1 Confers Biotic and Abiotic Stress Tolerance in Rice. PLoS One. 2015:10(6). 10.1371/JOURNAL.PONE.0127831

Jores T, Tonnies J, Wrightsman T, Buckler ES, Cuperus JT, Fields S, and Queitsch C. Synthetic promoter designs enabled by a comprehensive analysis of plant core promoters. Nature Plants 2021 7:6. 2021:7(6):842–855. 10.1038/s41477-021-00932-y

Jung SE, Bang SW, Kim SH, Seo JS, Yoon H Bin, Kim YS, and Kim JK. Overexpression of OsERF83, a Vascular Tissue-Specific Transcription Factor Gene, Confers Drought Tolerance in Rice. Int J Mol Sci. 2021:22(14). 10.3390/IJMS22147656

Junion G, Spivakov M, Girardot C, Braun M, Gustafson EH, Birney E, and Furlong EEM. A transcription factor collective defines cardiac cell fate and reflects lineage history. Cell. 2012:148(3):473–486. 10.1016/J.CELL.2012.01.030

Juven-Gershon T and Kadonaga JT. Regulation of gene expression via the core promoter and the basal transcriptional machinery. 2009. 10.1016/j.ydbio.2009.08.009

Kajala K, Covshoff S, Karki S, Woodfield H, Tolley BJ, Dionora MJA, Mogul RT, Mabilangan AE, Danila FR, Hibberd JM, et al. Strategies for engineering a two-celled C4 photosynthetic pathway into rice. J Exp Bot. 2011:62(9):3001–3010. 10.1093/JXB/ERR022

Kanaoka MM, Pillitteri LJ, Fujii H, Yoshida Y, Bogenschutz NL, Takabayashi J, Zhu JK, and Torii KU. SCREAM/ICE1 and SCREAM2 Specify Three Cell-State Transitional Steps Leading to Arabidopsis Stomatal Differentiation. Plant Cell. 2008:20(7):1775–1785. 10.1105/TPC.108.060848

Kirschner S, Woodfield H, Prusko K, Koczor M, Gowik U, Hibberd JM, and Westhoff P. Expression of SULTR2;2, encoding a low-affinity sulphur transporter, in the Arabidopsis bundle sheath and vein cells is mediated by a positive regulator. J Exp Bot. 2018:69(20):4897–4906. 10.1093/jxb/ery263

Krumlauf R. Hox genes in vertebrate development. Cell. 1994:78(2):191–201. 10.1016/0092-8674(94)90290-9

Kubo M, Udagawa M, Nishikubo N, Horiguchi G, Yamaguchi M, Ito J, Mimura T, Fukuda H, and Demura T. Transcription switches for protoxylem and metaxylem vessel formation. Genes Dev. 2005:19(16):1855–1860. 10.1101/GAD.1331305

Kulkarni MM and Arnosti DN. Information display by transcriptional enhancers. Development. 2003:130(26):6569–6575. 10.1242/DEV.00890

Kummari D, Palakolanu SR, Kishor PBK, Bhatnagar-Mathur P, Singam P, Vadez V, and Sharma KK. An update and perspectives on the use of promoters in plant genetic engineering. J Biosci. 2020:45(1):1–24. 10.1007/S12038-020-00087-6/TABLES/7

Kurt F, Filiz E, and Aydın A. Sulfite Reductase (SiR) Gene in Rice (Oryza sativa): Bioinformatics and Expression Analyses Under Salt and Drought Stresses. J Plant Growth Regul. 2022:41(6):2246–2260. 10.1007/S00344-021-10438-8/FIGURES/9

Lee DY, Hua L, Khoshravesh R, Giuliani R, Kumar I, Cousins A, Sage TL, Hibberd JM, and Brutnell TP. Engineering chloroplast development in rice through cell-specific control of endogenous genetic circuits. Plant Biotechnol J. 2021:19(11):2291–2303. 10.1111/PBI.13660

Leegood RC. Roles of the bundle sheath cells in leaves of C_3_ plants. J Exp Bot. 2008:59(7):1663–1673. 10.1093/jxb/erm335

Lewis EB. A gene complex controlling segmentation in Drosophila. Nature 1978 276:5688. 1978:276(5688):565–570. 10.1038/276565a0

Li E, Liu H, Huang L, Zhang X, Dong X, Song W, Zhao H, and Lai J. Long-range interactions between proximal and distal regulatory regions in maize. Nat Commun. 2019:10(1). 10.1038/S41467-019-10603-4

Li L, Zhu Q, He X, Sinha S, and Halfon MS. Large-scale analysis of transcriptional cis-regulatory modules reveals both common features and distinct subclasses. Genome Biol. 2007:8(6). 10.1186/GB-2007-8-6-R101

Lin R, Zhao W, Meng X, and Peng YL. Molecular cloning and characterization of a rice gene encoding AP2/EREBP-type transcription factor and its expression in response to infection with blast fungus and abiotic stresses. Physiol Mol Plant Pathol. 2007:70(1–3):60–68. 10.1016/J.PMPP.2007.06.002

Liu L, Adrian J, Pankin A, Hu J, Dong X, Von Korff M, and Turck F. Induced and natural variation of promoter length modulates the photoperiodic response of FLOWERING LOCUS T. Nat Commun. 2014:5. 10.1038/ncomms5558

Liu L, Du Y, Shen X, Li M, Sun W, Huang J, Liu Z, Tao Y, Zheng Y, Yan J, et al. KRN4 Controls Quantitative Variation in Maize Kernel Row Number. PLoS Genet. 2015:11(11):e1005670. 10.1371/JOURNAL.PGEN.1005670

MacAlister CA, Ohashi-Ito K, and Bergmann DC. Transcription factor control of asymmetric cell divisions that establish the stomatal lineage. Nature 2006 445:7127. 2006:445(7127):537–540. 10.1038/nature05491

Makino A, Sakuma H, Sudo E, and Mae T. Differences between Maize and Rice in N-use Efficiency for Photosynthesis and Protein Allocation.

Maruyama-Nakashita A, Nakamura Y, Tohge T, Saito K, and Takahashi H. Arabidopsis SLIM1 is a central transcriptional regulator of plant sulfur response and metabolism. Plant Cell. 2006:18(11):3235–3251. 10.1105/tpc.106.046458

McClung CR. Plant circadian rhythms. Plant Cell. 2006:18(4):792–803. 10.1105/TPC.106.040980

Mitchell PL and Sheehy JE. Supercharging rice photosynthesis to increase yield. New Phytol. 2006:171(4):688–693. 10.1111/J.1469-8137.2006.01855.X

Miyashima S, Roszak P, Sevilem I, Toyokura K, Blob B, Heo J ok, Mellor N, Help-Rinta-Rahko H, Otero S, Smet W, et al. Mobile PEAR transcription factors integrate positional cues to prime cambial growth. Nature. 2019:565(7740):490–494. 10.1038/S41586-018-0839-Y

Moreno-Risueno MA, Sozzani R, Yardimci GG, Petricka JJ, Vernoux T, Blilou I, Alonso J, Winter CM, Ohler U, Scheres B, et al. Transcriptional control of tissue formation throughout root development. Science (1979). 2015:350(6259):426–430. 10.1126/SCIENCE.AAD1171/SUPPL_FILE/TABLE_S6.XLSX

Nagel DH and Kay SA. Complexity in the wiring and regulation of plant circadian networks. Curr Biol. 2012:22(16). 10.1016/J.CUB.2012.07.025

Nakashima K, Ito Y, and Yamaguchi-Shinozaki K. Transcriptional Regulatory Networks in Response to Abiotic Stresses in Arabidopsis and Grasses. Plant Physiol. 2009:149(1):88. 10.1104/PP.108.129791

Nomura M, Higuchi T, Ishida Y, Ohta S, Komari T, Imaizumi N, Miyao-Tokutomi M, Matsuoka M, and Tajima S. Differential expression pattern of C4 bundle sheath expression genes in rice, a C3 plant. Plant Cell Physiol. 2005a:46(5):754–761. 10.1093/PCP/PCI078

Nomura M, Higuchi T, Ishida Y, Ohta S, Komari T, Imaizumi N, Miyao-Tokutomi M, Matsuoka M, and Tajima S. Differential expression pattern of C4 bundle sheath expression genes in rice, a C3 plant. Plant Cell Physiol. 2005b:46(5):754–761. 10.1093/PCP/PCI078

Ohashi-Ito K and Bergmann DC. Arabidopsis FAMA Controls the Final Proliferation/Differentiation Switch during Stomatal Development. Plant Cell. 2006:18(10):2493–2505. 10.1105/TPC.106.046136

Page MT, Parry MAJ, and Carmo-Silva E. A high-throughput transient expression system for rice. Plant Cell Environ. 2019:42(7):2057–2064. 10.1111/PCE.13542

Panne D. The enhanceosome. Curr Opin Struct Biol. 2008:18(2):236–242. 10.1016/J.SBI.2007.12.002

Pillitteri LJ, Sloan DB, Bogenschutz NL, and Torii KU. Termination of asymmetric cell division and differentiation of stomata. Nature 2006 445:7127. 2006:445(7127):501–505. 10.1038/nature05467

Qu X, Zou J, Wang J, Yang K, Wang X, and Le J. A Rice R2R3-Type MYB Transcription Factor OsFLP Positively Regulates Drought Stress Response via OsNAC. Int J Mol Sci. 2022:23(11). 10.3390/IJMS23115873

Que Q, Wang HY, English JJ, and Jorgensen RA. The Frequency and Degree of Cosuppression by Sense Chalcone Synthase Transgenes Are Dependent on Transgene Promoter Strength and Are Reduced by Premature Nonsense Codons in the Transgene Coding Sequence. Plant Cell. 1997:9(8):1357. 10.1105/TPC.9.8.1357

Ray-Jones H and Spivakov M. Transcriptional enhancers and their communication with gene promoters. Cellular and Molecular Life Sciences. 2021:78(19–20):6453. 10.1007/S00018-021-03903-W

Reyna-Llorens I, Burgess SJ, Reeves G, Singh P, Stevenson SR, Williams BP, Stanley S, and Hibberd JM. Ancient duons may underpin spatial patterning of gene expression in C4 leaves. Proc Natl Acad Sci U S A. 2018:115(8):1931–1936. 10.1073/PNAS.1720576115/SUPPL_FILE/PNAS.1720576115.SD07.XLSX

Rose AB. Introns as Gene Regulators: A Brick on the Accelerator. Front Genet. 2019:9(FEB). 10.3389/FGENE.2018.00672

Rose AB, Elfersi T, Parra G, and Korf I. Promoter-proximal introns in Arabidopsis thaliana are enriched in dispersed signals that elevate gene expression. Plant Cell. 2008:20(3):543–551. 10.1105/TPC.107.057190

Sage RF. The evolution of C_4_ photosynthesis. New Phytologist. 2004:161(2):341–370. 10.1111/j.1469-8137.2004.00974.x

Schmitz RJ, Grotewold E, and Stam M. Cis-regulatory sequences in plants: Their importance, discovery, and future challenges. Plant Cell. 2022:34(2):718–741. 10.1093/PLCELL/KOAB281

Shimizu H, Sato K, Berberich T, Miyazaki A, Ozaki R, Imai R, and Kusano T. LIP19, a basic region leucine zipper protein, is a Fos-like molecular switch in the cold signaling of rice plants. Plant Cell Physiol. 2005:46(10):1623–1634. 10.1093/PCP/PCI178

Shlyueva D, Stampfel G, and Stark A. Transcriptional enhancers: from properties to genome-wide predictions. Nat Rev Genet. 2014:15(4):272–286. 10.1038/NRG3682

Smale ST and Kadonaga JT. The RNA polymerase II core promoter. Annu Rev Biochem. 2003:72(Volume 72, 2003):449–479. 10.1146/ANNUREV.BIOCHEM.72.121801.161520/CITE/REFWORKS

Sparks EE, Drapek C, Gaudinier A, Li S, Ansariola M, Shen N, Hennacy JH, Zhang J, Turco G, Petricka JJ, et al. Establishment of Expression in the SHORTROOT-SCARECROW Transcriptional Cascade through Opposing Activities of Both Activators and Repressors. Dev Cell. 2016:39(5):585–596. 10.1016/j.devcel.2016.09.031

Spitz F and Furlong EEM. Transcription factors: from enhancer binding to developmental control. Nat Rev Genet. 2012:13(9):613–626. 10.1038/NRG3207

Stam M, Belele C, Dorweiler JE, and Chandler VL. Differential chromatin structure within a tandem array 100 kb upstream of the maize b1 locus is associated with paramutation. Genes Dev. 2002:16(15):1906–1918. 10.1101/GAD.1006702

Stergachis AB, Haugen E, Shafer A, Fu W, Vernot B, Reynolds A, Raubitschek A, Ziegler S, LeProust EM, Akey JM, et al. Exonic transcription factor binding directs codon choice and impacts protein evolution. Science. 2013:342(6164):1367. 10.1126/SCIENCE.1243490

Studer A, Zhao Q, Ross-Ibarra J, and Doebley J. Identification of a functional transposon insertion in the maize domestication gene tb1. Nat Genet. 2011:43(11):1160. 10.1038/NG.942

Swift J, Luginbuehl LH, Schreier TB, Donald RM, Lee TA, Nery JR, Ecker JR, and Hibberd JM. Single nuclei sequencing reveals C4 photosynthesis is based on rewiring of ancestral cell identity networks. bioRxiv. 2023:2023.10.26.562893. 10.1101/2023.10.26.562893

Tezuka D, Kawamata A, Kato H, Saburi W, Mori H, and Imai R. The rice ethylene response factor OsERF83 positively regulates disease resistance to Magnaporthe oryzae. Plant Physiol Biochem. 2019:135:263–271. 10.1016/J.PLAPHY.2018.12.017

Thanos D and Maniatis T. Virus induction of human IFNβ gene expression requires the assembly of an enhanceosome. Cell. 1995:83(7):1091–1100. 10.1016/0092-8674(95)90136-1

Tsuda K and Somssich IE. Transcriptional networks in plant immunity. New Phytol. 2015:206(3):932–947. 10.1111/NPH.13286

Verma V, Ravindran P, and Kumar PP. Plant hormone-mediated regulation of stress responses. BMC Plant Biol. 2016:16(1):1–10. 10.1186/S12870-016-0771-Y/FIGURES/1

Wang L, Wan MC, Liao RY, Xu J, Xu ZG, Xue HC, Mai YX, and Wang JW. The maturation and aging trajectory of Marchantia polymorpha at single-cell resolution. Dev Cell. 2023:58(15):1429–1444.e6. 10.1016/J.DEVCEL.2023.05.014

Wang Y, Huan Q, Li K, and Qian W. Single-cell transcriptome atlas of the leaf and root of rice seedlings. Journal of Genetics and Genomics. 2021:48(10):881–898. 10.1016/J.JGG.2021.06.001

Weber B, Zicola J, Oka R, and Stam M. Plant Enhancers: A Call for Discovery. Trends Plant Sci. 2016:21(11):974–987. 10.1016/J.TPLANTS.2016.07.013

Wiludda C, Schulze S, Gowik U, Engelmann S, Koczor M, Streubel M, Bauwe H, and Westhoff P. Regulation of the Photorespiratory GLDPA Gene in C4 Flaveria: An Intricate Interplay of Transcriptional and Posttranscriptional Processes. Plant Cell. 2012:24(1):137. 10.1105/TPC.111.093872

Xiong H, Hua L, Reyna-Llorens I, Shi Y, Chen KM, Smirnoff N, Kromdijk J, and Hibberd JM. Photosynthesis-independent production of reactive oxygen species in the rice bundle sheath during high light is mediated by NADPH oxidase. Proc Natl Acad Sci U S A. 2021:118(25):e2022702118. 10.1073/PNAS.2022702118/SUPPL_FILE/PNAS.2022702118.SD04.XLSX

Yaish MW, El-Kereamy A, Zhu T, Beatty PH, Good AG, Bi YM, and Rothstein SJ. The APETALA-2-like transcription factor OsAP2-39 controls key interactions between abscisic acid and gibberellin in rice. PLoS Genet. 2010:6(9). 10.1371/JOURNAL.PGEN.1001098

Yamamoto Y, Ichida H, Abe T, Suzuki Y, Sugano S, and Obokata J. Differentiation of core promoter architecture between plants and mammals revealed by LDSS analysis. Nucleic Acids Res. 2007a:35(18):6219. 10.1093/NAR/GKM685

Yamamoto YY, Ichida H, Matsui M, Obokata J, Sakurai T, Satou M, Seki M, Shinozaki K, and Abe T. Identification of plant promoter constituents by analysis of local distribution of short sequences. BMC Genomics. 2007b:8(1):1–23. 10.1186/1471-2164-8-67/FIGURES/9

Yan W, Chen D, Schumacher J, Durantini D, Engelhorn J, Chen M, Carles CC, and Kaufmann K. Dynamic control of enhancer activity drives stage-specific gene expression during flower morphogenesis. Nat Commun. 2019:10(1). 10.1038/S41467-019-09513-2

Zhang C, Ding Z, Wu K, Yang L, Li Y, Yang Z, Shi S, Liu X, Zhao S, Yang Z, et al. Suppression of Jasmonic Acid-Mediated Defense by Viral-Inducible MicroRNA319 Facilitates Virus Infection in Rice. Mol Plant. 2016:9(9):1302–1314. 10.1016/J.MOLP.2016.06.014

Zhang C, Zhang J, Liu H, Qu X, Wang J, He Q, Zou J, Yang K, and Le J. Transcriptomic analysis reveals the role of FOUR LIPS in response to salt stress in rice. Plant Mol Biol. 2022:110(1–2):37–52. 10.1007/S11103-022-01282-9/FIGURES/7

Zhang H, Zhang J, Quan R, Pan X, Wan L, and Huang R. EAR motif mutation of rice OsERF3 alters the regulation of ethylene biosynthesis and drought tolerance. Planta. 2013:237(6):1443–1451. 10.1007/S00425-013-1852-X

Zhang X, Bao Y, Shan D, Wang Z, Song X, Wang Z, Wang J, He L, Wu L, Zhang Z, et al. Magnaporthe oryzae Induces the Expression of a MicroRNA to Suppress the Immune Response in Rice. Plant Physiol. 2018:177(1):352. 10.1104/PP.17.01665

Zhao H, Yang M, Bishop J, Teng Y, Cao Y, Beall BD, Li S, Liu T, Fang Q, Fang C, et al. Identification and functional validation of super-enhancers in Arabidopsis thaliana. Proc Natl Acad Sci U S A. 2022:119(48):e2215328119. 10.1073/PNAS.2215328119/SUPPL_FILE/PNAS.2215328119.SD02.XLSX

