## Supplementary material for "A transcription factor quintet orchestrating bundle sheath expression in rice": S Figures

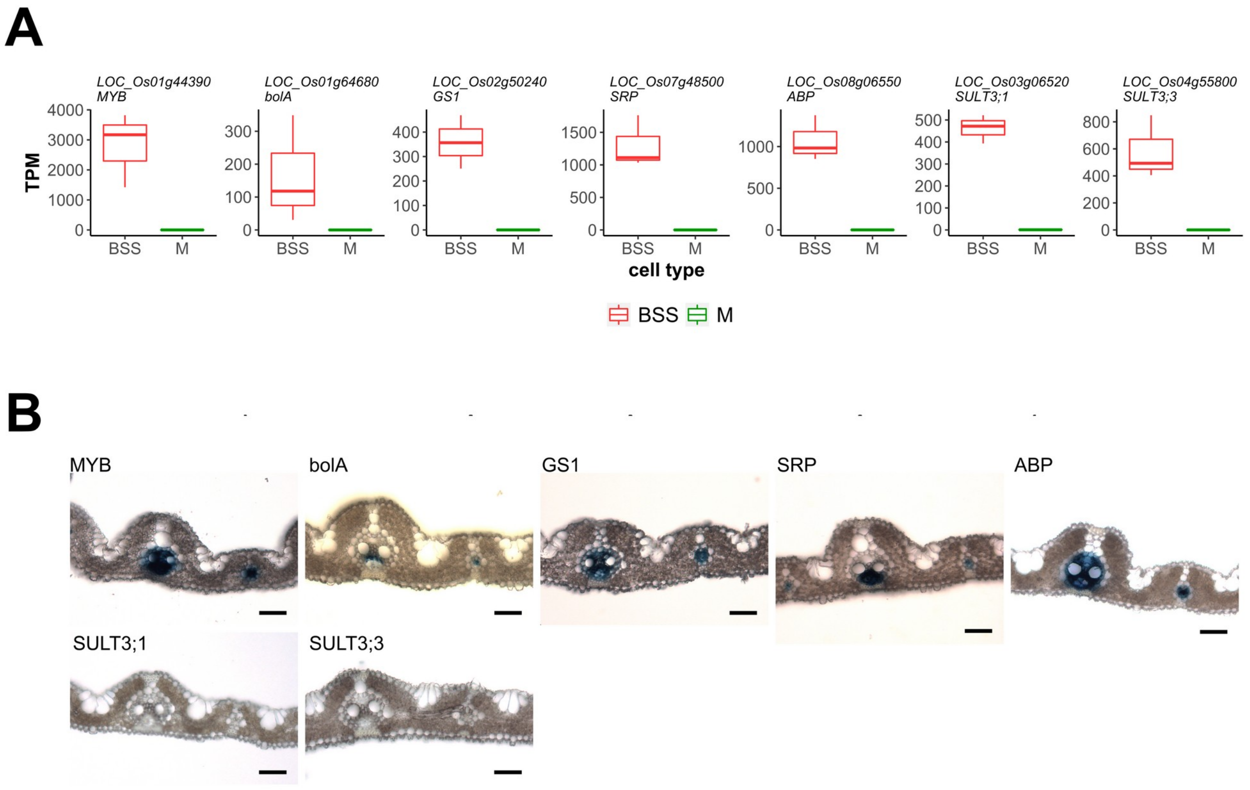


C

| NO. | GENE ID | MSU7 annotation | Symbol | Region cloned | GUS localisation |
|---|---|---|---|---|---|
| 1 | LOC_Os01g44390 | MYB family transcription factor | MYB | -1641-+178 | V |
| 2 | LOC_Os01g64680 | bolA | bolA | -960-+345 | V |
| 3 | LOC_Os02g50240 | glutamine synthetase | GS1 | -1976-+804 | V |
| 4 | LOC_Os07g48500 | stress responsive protein | SRP | -1373-+496 | V |
| 5 | LOC_Os08g06550 | acyl CoA binding protein | ABP | -1468-+805 | V |
| 6 | LOC_Os03g06520 | sulfate transporter | SULT3;1 | -1508-+795 | No expression |
| 7 | LOC_Os04g55800 | sulfate transporter | SULT3;3 | -2052-+264 | No expression |

**Supplemental Figure 1. Analysis of seven rice promoters identified after analysis of transcripts that accumulate preferentially in bundle sheath strands. (A)** Transcript abundance (Transcript per million, TPM) derived from each gene. **(B)** Representative GUS staining images using promoters from *MYB, bolA, GS1, SRP, ABP, SULT3;1* and *SULT3;3*. Scale bars represent 50 µm. **(C)** Summary of promoters tested including gene information, upstream flanking region (relative to translational start site), and GUS localisation. Abbreviations: BSS, Bundle Sheath Strands (vascular bundle and bundle sheath); M, Mesophyll; V, vascular bundle (cell types including xylem, phloem which are surrounded by bundle sheath cells).

A

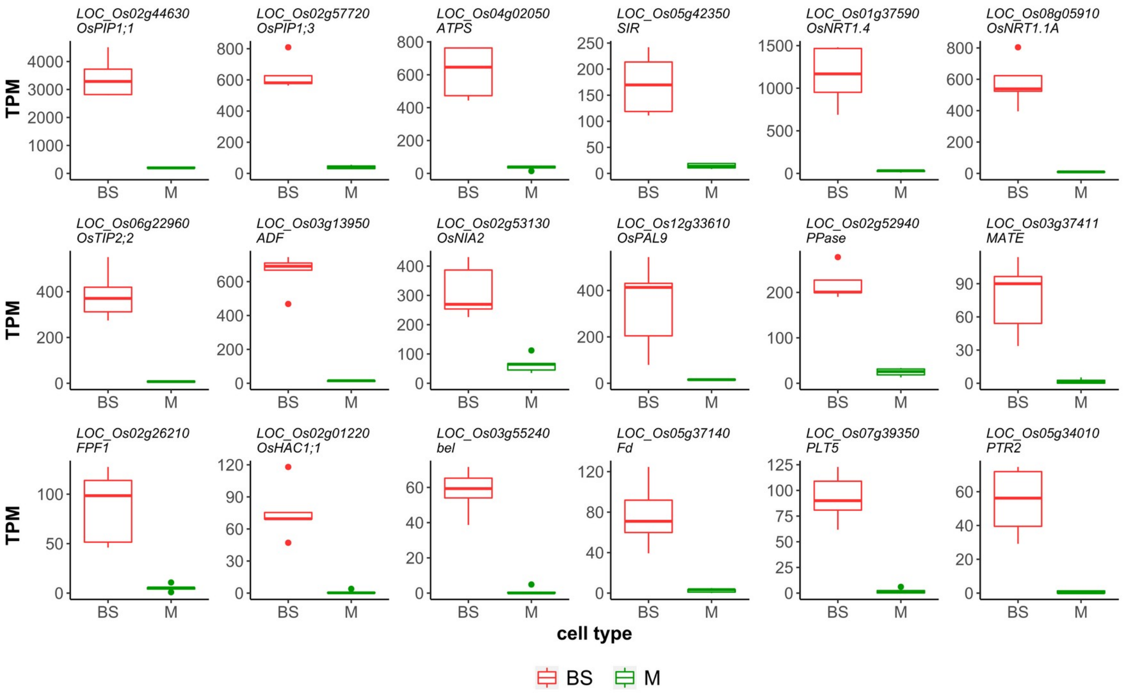


B

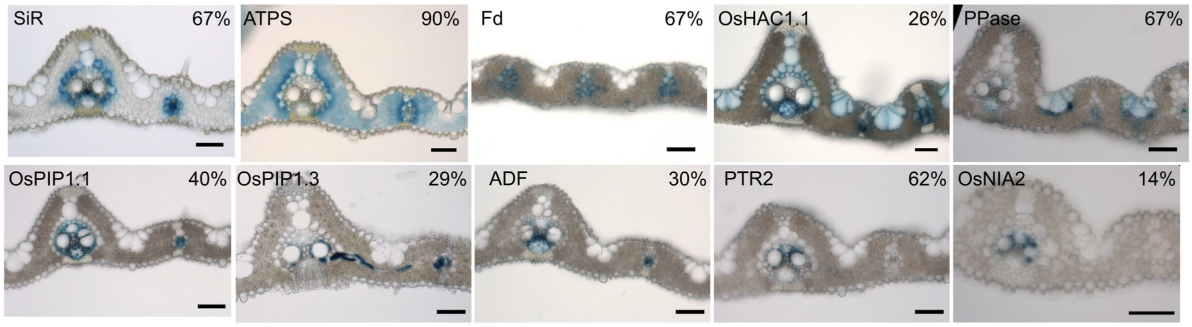


C

| NO. | GENE ID | MSU7 annotation | Symbol | Region cloned | GUS localisation |
|---|---|---|---|---|---|
| 1 | LOC_Os04g02050 | bifunctional 3-phosphoadenosine 5-phosphosulfate synthetase | ATPS | -1769 - +2081 | BS and M |
| 2 | LOC_Os05g42350 | ferredoxin--nitrite reductase | SIR | -2571 - +42 | BS and V |
| 3 | LOC_Os05g37140 | 2Fe-2S iron-sulfur cluster binding domain containing protein (ferredoxin) | Fd | -1745 - +108 | BS, M and V |
| 4 | LOC_Os02g01220 | rhodanese-like domain containing protein | OsHAC1;1 | -3011 - -1 | BS, V and epidermis |
| 5 | LOC_Os02g52940 | soluble inorganic pyrophosphatase | PPase | -1456 - +9 | V |
| 6 | LOC_Os02g44630 | aquaporin protein | OsPIP1;1 | -3079 - -1 | V |
| 7 | LOC_Os02g57720 | aquaporin protein | OsPIP1;3 | -2364 - -1 | V |
| 8 | LOC_Os03g13950 | actin-depolymerizing factor | ADF | -2937 - +174 | V |
| 9 | LOC_Os05g34010 | peptide transporter PTR2 | PTR2 | -2783 - +27 | V |
| 10 | LOC_Os02g53130 | nitrate reductase | OsNIA2 | -3191 - +165 | V |
| 11 | LOC_Os12g33610 | phenylalanine ammonia-lyase | OsPAL9 | -2631 - +117 | No expression |
| 12 | LOC_Os01g37590 | peptide transporter PTR2 | OsNRT1.4 | -2576 - +147 | No expression |
| 13 | LOC_Os03g55240 | cytochrome P450 | bel | -2700 - +210 | No expression |
| 14 | LOC_Os06g22960 | aquaporin protein | OsTIP2;2 | -2984 - -1 | No expression |
| 15 | LOC_Os08g05910 | peptide transporter PTR2 | OsNRT1.1A | -1818 - +249 | No expression |
| 16 | LOC_Os07g39350 | transporter family protein | PLT5 | -2520 - +87 | No expression |
| 17 | LOC_Os03g37411 | MATE efflux family protein | MATE | -2904 - +198 | No expression |
| 18 | LOC_Os02g26210 | flowering promoting factor-like 1 | FPF1 | -2464 - +162 | No expression |

**Supplemental Figure 2. Identification of bundle sheath specific promoters after analysis of transcripts that accumulate preferentially in bundle sheath compared with mesophyll cells.** (A) Transcript abundance (Transcript per million, TPM) of eighteen candidate genes in bundle sheath (BS) and mesophyll (M) cells. (B) Representative cross sections of leaves after GUS staining showing bundle sheath or vascular bundle expression. Percentage of transgenic lines with each GUS pattern indicated upper-right, in the remaining lines GUS was not detectable. Scale bars represent 50 µm. (C) Summary of gene information, region cloned (relative to translational start site) and GUS localisation. Abbreviations: BS, Bundle Sheath; M, Mesophyll; V, Vascular bundle (cell types including xylem, phloem which are surrounded by bundle sheath cells).

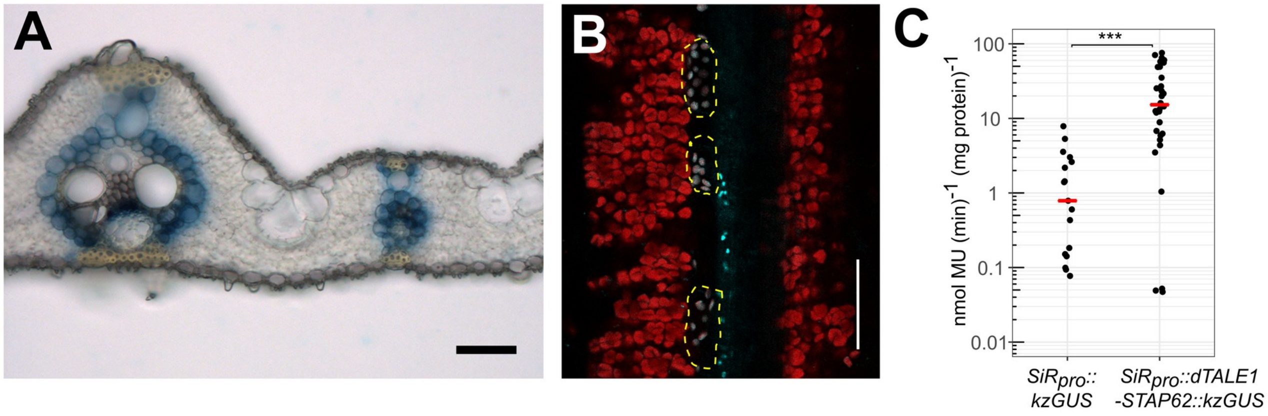


**Supplemental Figure 3. The domesticated *SiR* promoter combined with the dTALE/STAP system drives strong expression in the rice bundle sheath. (A)** Representative image of expression from the domesticated *SiR* promoter driving dTALE and STAP4-GUS in transverse section of rice leaf. (**B**) Representative image of expression from of nuclear localized mTurquoise2 fluorescent protein under control of the domesticated *SiR* promoter driving dTALE, and mTurquoise2 downstream of STAP62. Bundle sheath cells marked by yellow dashed lines, and red indicates chlorophyll autofluorescence. **(C)** GUS activity mediated by dTALE and STAP62 compared with the native *SiR* promoter. Data were subjected to a Wilcoxon test, *** represents $P < 0.001$.

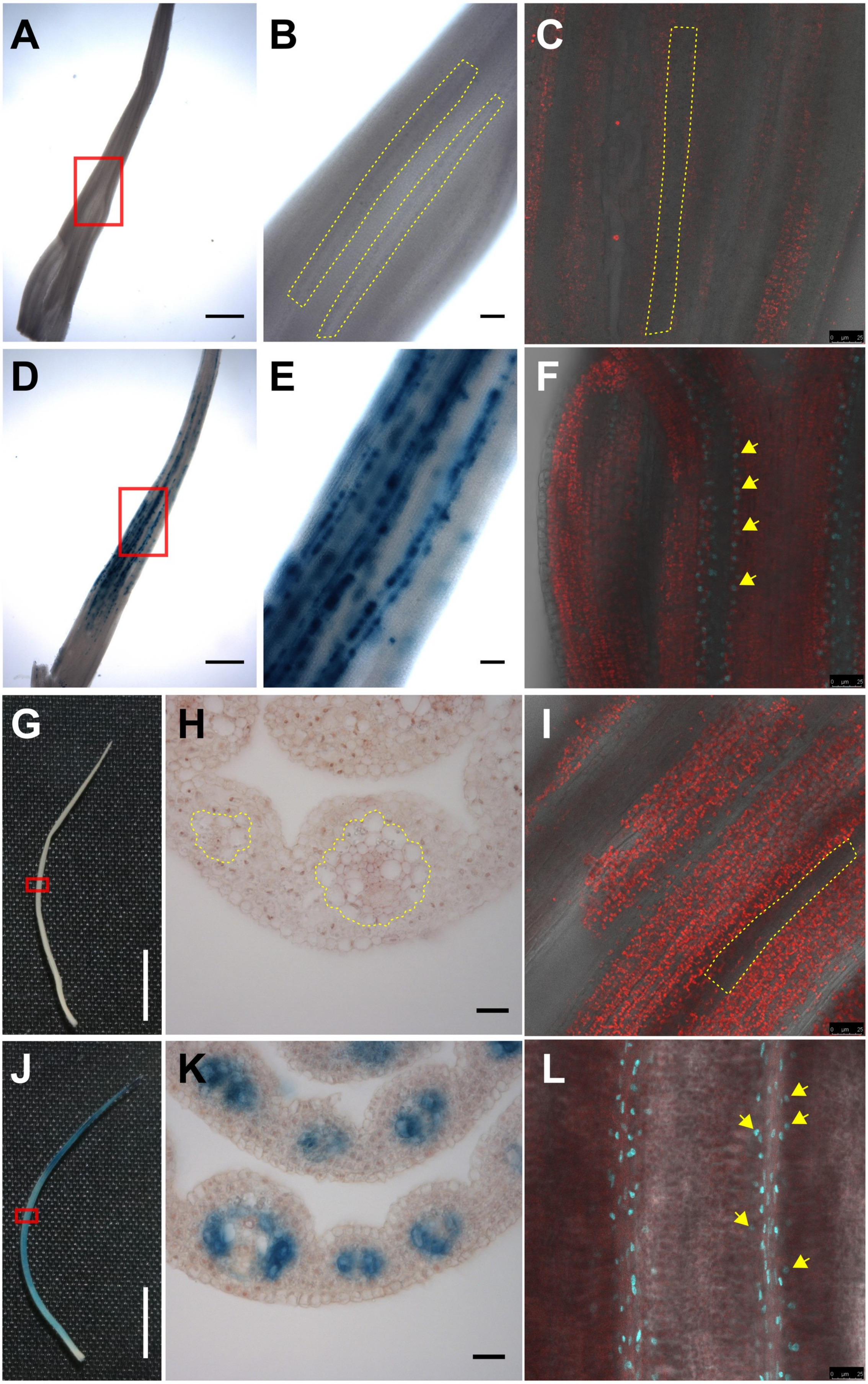
A
B
C
D
E
F
G
H
I
J
K
L

**Supplemental Figure 4. The rice *SiR* promoter drives expression in bundle sheath cells earlier than the *Zoysia japonica PCK* promoter.** GUS in transgenic rice plants expressing *ZjPCKpro::kzGUS* (A, B, G, H) and *SiRpro::kzGUS* (**D, E, J, K**) from 5-mm (**A, B, D, E**) and 2-cm (**G, H, J, K**) fourth leaves. **B&E** show magnified views of the boxed regions in **A** and **D**. Expression of green fluorescent protein (**C, I**) and mTurquoise2 (**F, L**) in transgenic plants expressing *ZjPCKpro::dTALE1-STAP4::GFP-NLS* (**C,** I) and *SiRpro::H2B-mTurquoise2* (**F, L**) from 5-mm (**C, F**) and 2-cm (**I, L**) fourth leaves. NLS and H2B are nuclear localisation signal peptides. Yellow arrows indicate nuclei of bundle sheath cells expressing H2B-mTurquoise2, bundle sheath strands in *ZjPCKpro* lines were highlighted with yellow dash lines. Chlorophyll autofluorescence indicated in red (**C, F, I, L**). Scale bars represent 500 µm for **A, D**; 50 µm for **B, E, H, K**; 25 µm for **C, F, I, L**; 5 mm for **G, J.**

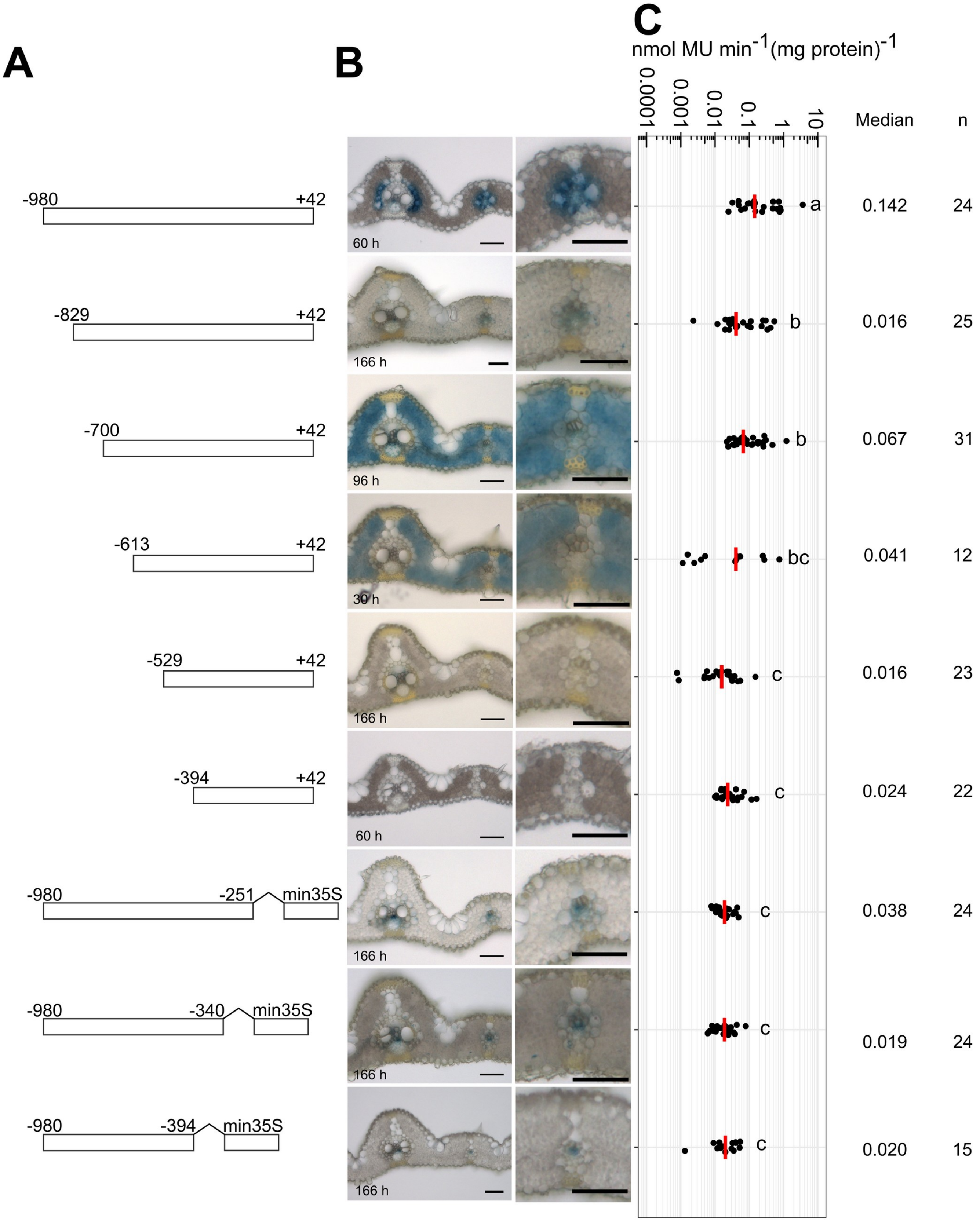


**Supplemental Figure 5. Impact of 5' and 3' deletions on patterning of GUS from the *SiR* promoter.** (A) Schematics showing sequences fused to GUS reporter. (**B**) Representative images of leaf cross sections after GUS staining. (**C**) Promoter activity determined by the fluorometric 4-methylumbelliferyl-β-D-glucuronide (MUG) assay. Data subjected to pairwise Wilcoxon test with Benjamini-Hochberg correction. Lines with differences in activity that were statistically significant (adjusted $P<0.05$) labelled with different letters. Median catalytic rate of GUS indicated with red line, n indicates total number of transgenic lines assessed. n indicates total number of transgenic lines assessed for each construct.

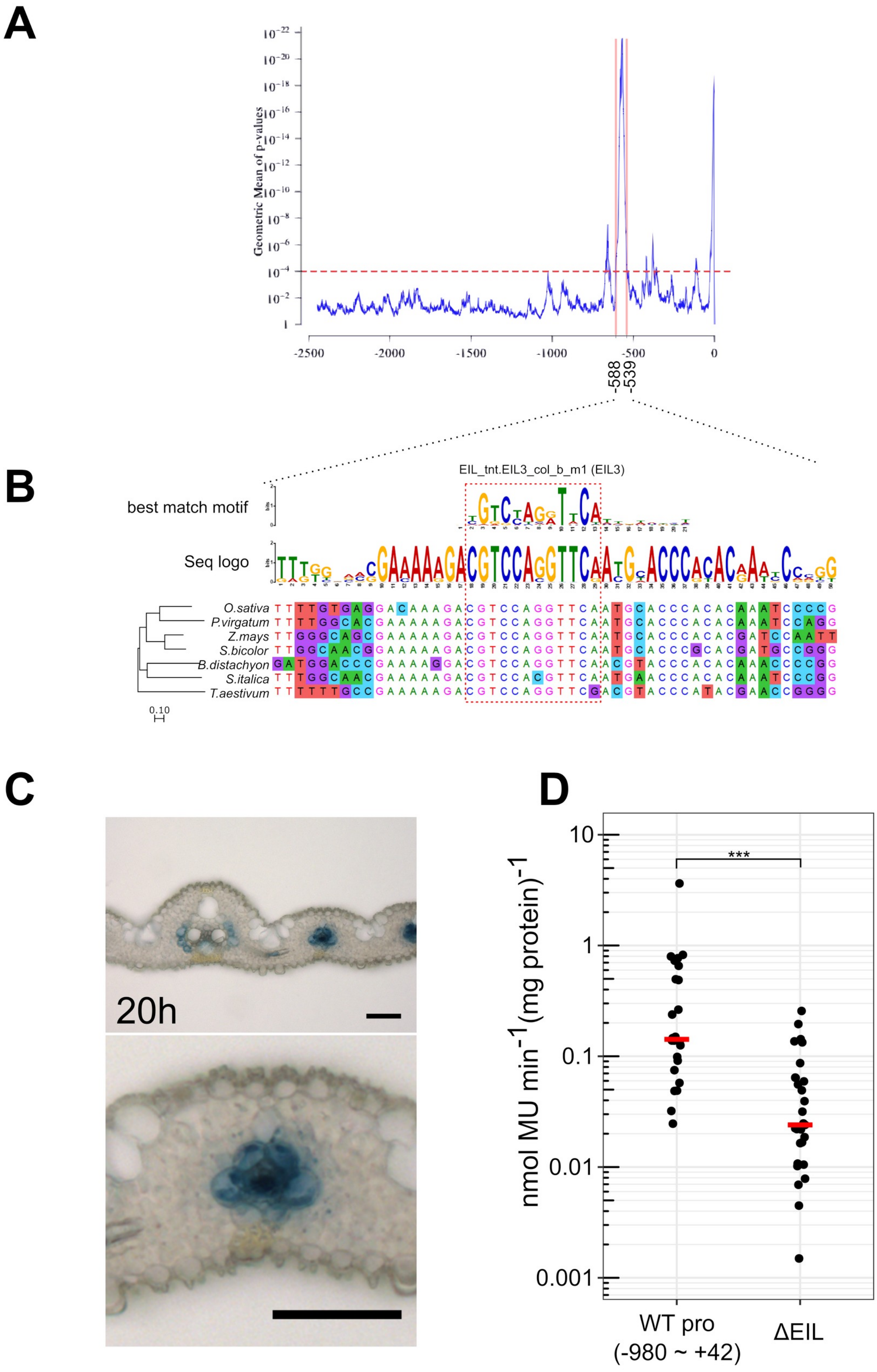


**Supplemental Figure 6. The evolutionally conserved *ETHYLENE INSENSITIVE3-LIKE* (*EIL*) binding site regulates expression level but not cell specificity of the *SiR* promoter.** (**A**) Conservation profile of 50-bp sliding window of the *SiR* promoter from seven grass species. (**B**) Sequence alignment at region -558 to -538 bp and the best match motif for this region. (**C**) Representative image of leaf cross sections after GUS staining $T_0$ transgenics transformed with deleted EIL site. (**D**) Promoter activity determined by the fluorometric 4-methylumbelliferyl-β-D-glucuronide (MUG) assay. Data were subjected to a Wilcoxon test, *** represents *P <0.001*.

**A**

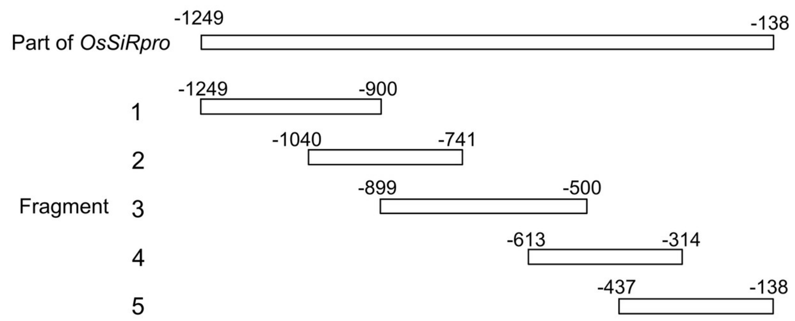


**B**

| Fragment | nucleotides | confidence | MSU ID | Symbol | TF family |
|---|---|---|---|---|---|
| 1 | -1249 .. -900 | D | LOC_Os07g05720 | OsTCP21 | TCP |
| 1 | -1249 .. -900 | D | LOC_Os12g37410 | OsOBF1,OsbZIP87 | bZIP_A/Ocs |
| 2 | -1040 .. -741 | D | LOC_Os12g37410 | OsOBF1,OsbZIP87 | bZIP_A/Ocs |
| 2 | -1040.. -741 | A | LOC_Os02g54160 | EREBP1 | ERF/DREB_A |
| 2 | -1040.. -741 | D | LOC_Os03g64260 | OsERF83 | ERF/DREB_A |
| 2 | -1040.. -741 | A | LOC_Os04g52090 | OsAP2-39 | ERF/DREB_A |
| 2 | -1040.. -741 | A | LOC_Os01g58420 | AP37,OsERF3 | ERF/DREB_A |
| 2 | -1040.. -741 | A | LOC_Os05g41780 | OsERF74 | ERF/DREB_A |
| 2 | -1040.. -741 | D | LOC_Os09g26420 | OsERF72 | ERF/DREB_A |
| 3 | -899 .. -500 | D | LOC_Os04g52090 | OsAP2-39 | ERF/DREB_A |
| 3 | -899 .. -500 | A | LOC_Os03g20780 | OsEIL1 | EIL |
| 3 | -899 .. -500 | A | LOC_Os07g48630 | OsEIL2 | EIL |
| 3 | -899 .. -500 | D | LOC_Os09g31400 | OsEIL3 | EIL |
| 4 | -613 .. -314 | A | LOC_Os01g69980 | TCP20 | TCP |
| 4 | -613 .. -314 | D | LOC_Os07g43420 | OsFLP | MYB_C |
| 4 | -613 .. -314 | B | LOC_Os08g43160 | PCF2,OsPCF2 | TCP |
| 5 | -437 .. -138 | D | LOC_Os01g55750 | OsTCP5 | TCP |
| 5 | -437 .. -138 | A | LOC_Os01g69980 | TCP20 | TCP |
| 5 | -437 .. -138 | D | LOC_Os03g21060 | NAC58, OsNAP | NAC_B |
| 5 | -437 .. -138 | D | LOC_Os07g05720 | OsTCP21 | TCP |
| 5 | -437 .. -138 | A | LOC_Os08g43160 | PCF2,OsPCF2 | TCP |

**C**

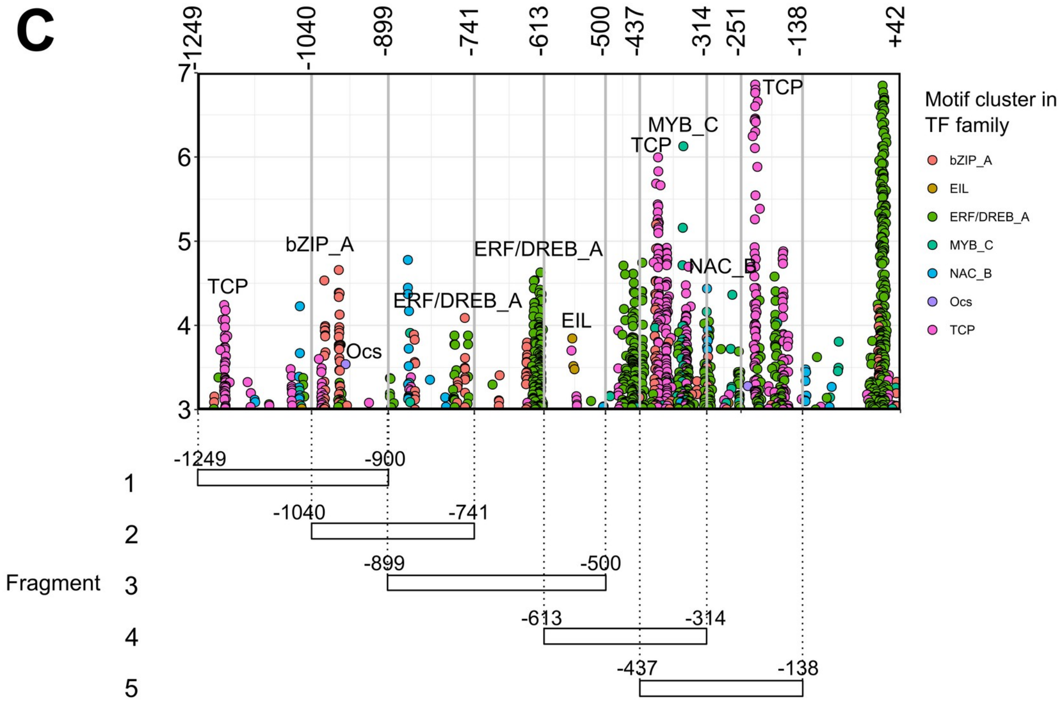

**D**

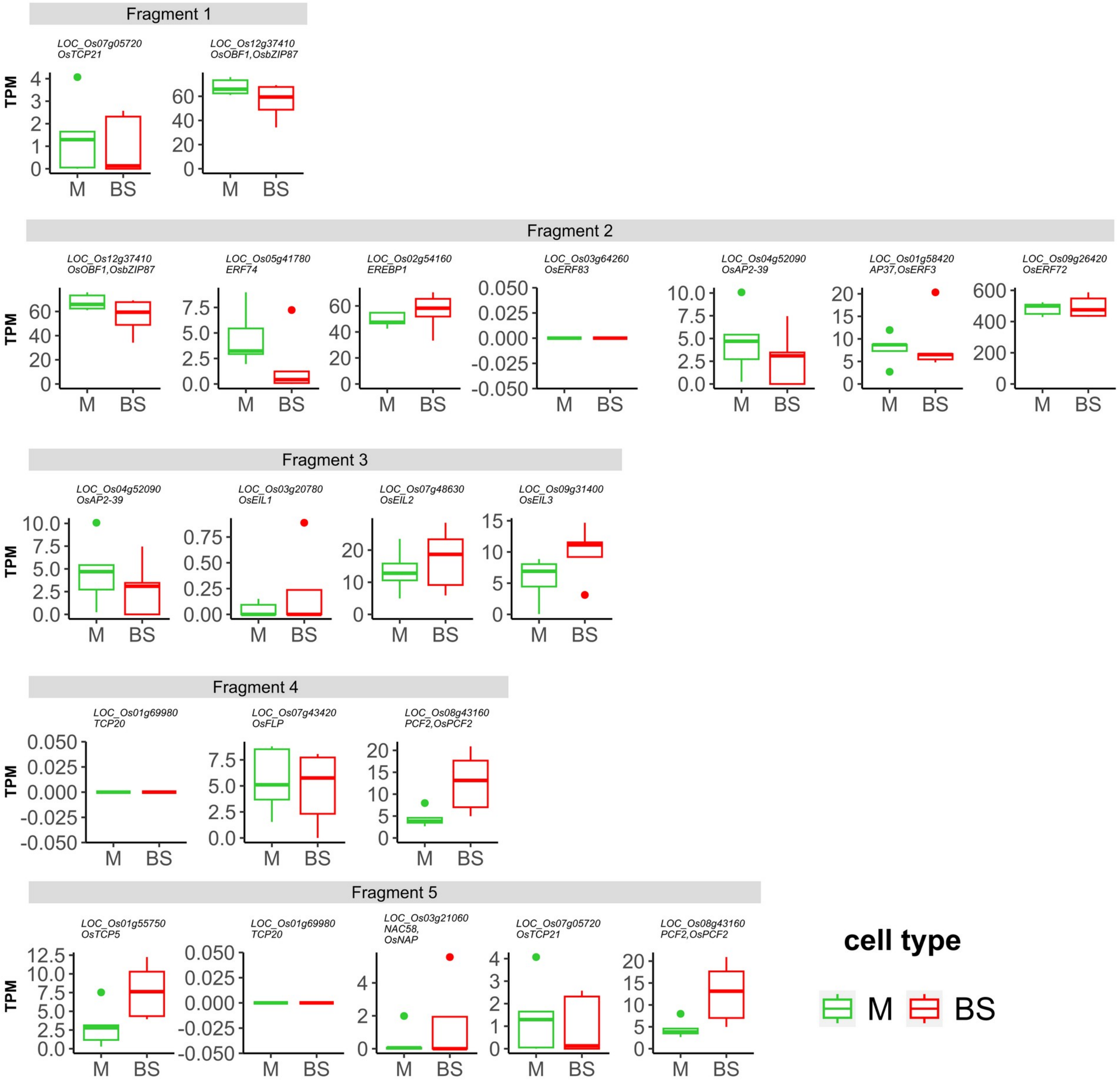


**Supplemental Figure 7. Identification of transcription factors interacting with the *SiR* promoter using Yeast one Hybridization*.*** (**A**) Schematic showing five fragments covering region of the *SiR* promoter used for yeast one hybrid analysis. (**B**) Transcription factors identified as interacting with each fragment. (**C**) Predicted binding sites of transcription factors in each fragment. (D) Transcript abundance for transcription factor genes in bundle sheath and mesophyll cells (Hua et al., 2021).

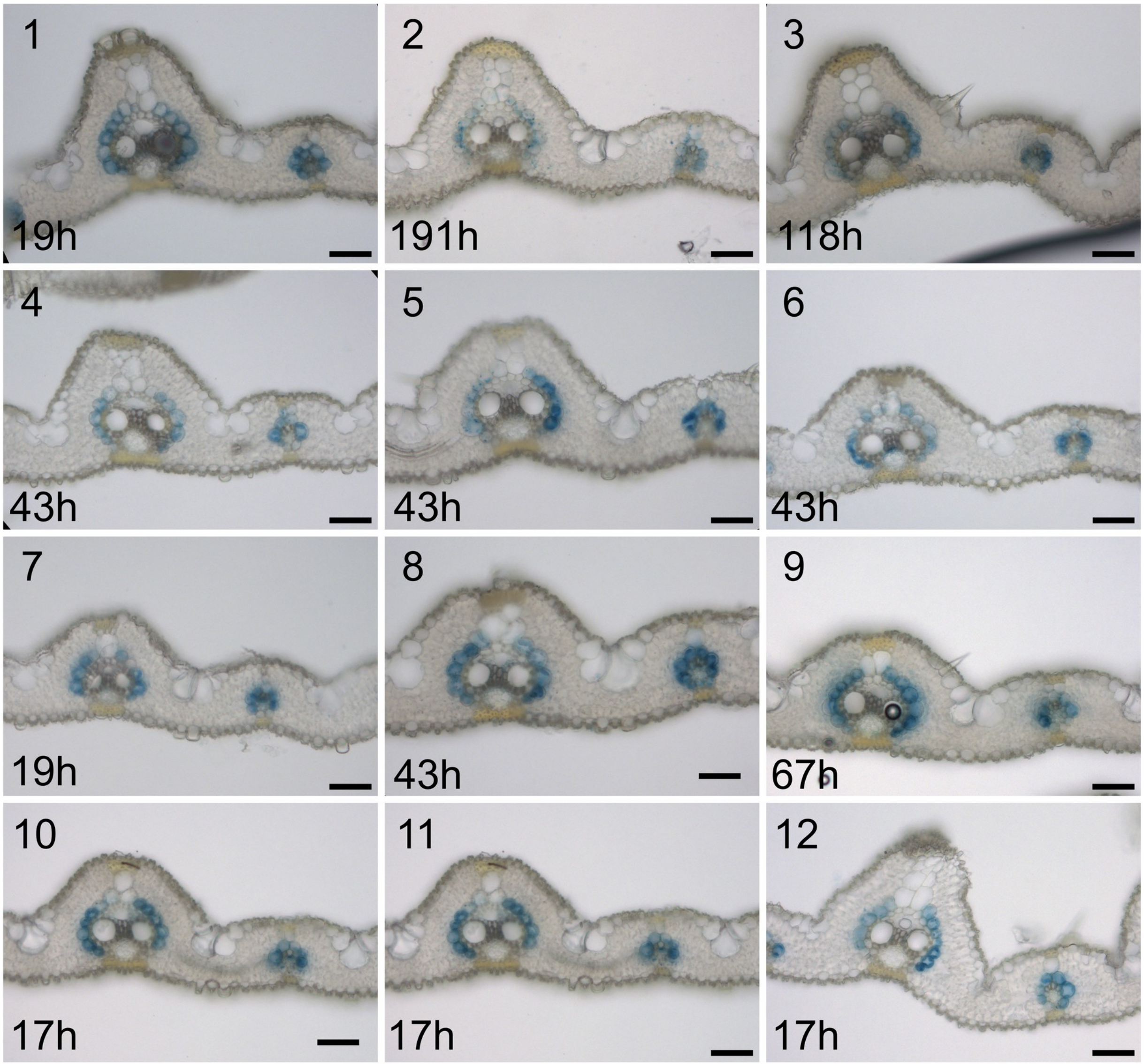


**Supplemental Figure 8. Nucleotides -980 to -829 in combination with nucleotides -251 to +42 produce bundle sheath specific expression.** Twelve independent lines assessed. The staining duration is displayed in the bottom-left corner, scale bars = 50 µm.

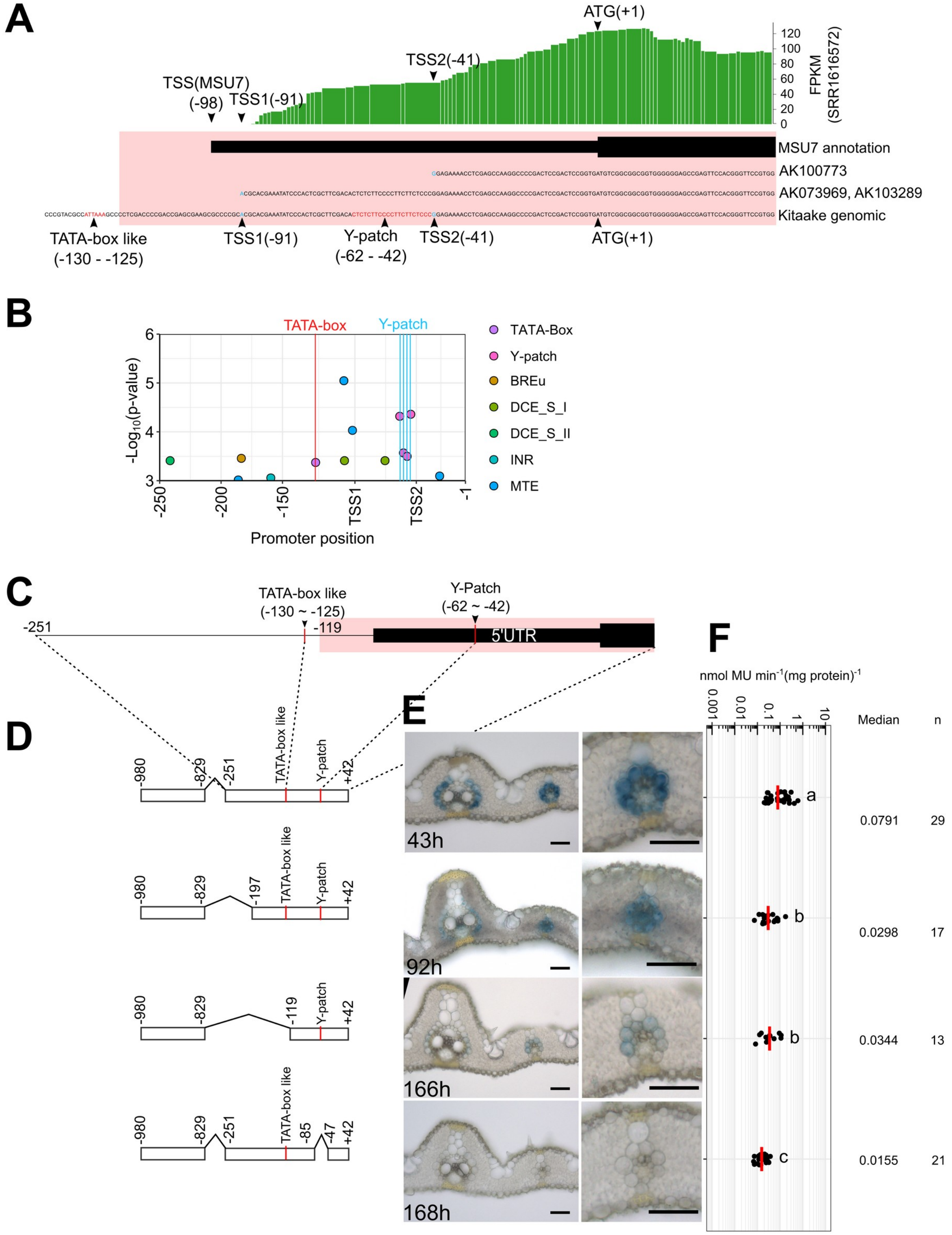


**Supplemental Figure 9. Nucleotides -251 to -1 serve as the core promoter.** (**A**) Two transcription start sites are supported by full length cDNA clones (TSS1: AK073969, AK103289; TSS2: AK100773) and leaf RNA sequencing reads (SRR1616572). Relative position to TATA box variant (TATA var. -130 to -125bp) and Y-Patch core promoter element (-62 to -42bp) depicted, and frequency of leaf RNAseq reads indicated with green bars. (**B**) Profile of general transcription factor binding site associated with RNA Polymerase II. (**C, D**) Diagrams showing position of core promoter elements TATA box and Y-patch between nucleotides -251 to +42 (**C**), and sequences fused to GUS (**D**). (**E**) Representative images of leaf cross sections after GUS staining with zoomed-in images of lateral veins shown in right panels, scale bars = 50 um. (**F**) Promoter activity determined by the fluorometric 4- methylumbelliferyl-β-D-glucuronide (MUG) assay. n indicates total number of transgenic lines assessed. Data subjected to pairwise Wilcoxon test with Benjamini-Hochberg correction. Lines with differences in activity that were statistically significant (adjusted $P<0.05$) labelled with different letters. Median catalytic rate of GUS indicated with red line, n indicates total number of transgenic lines assessed. Abbreviations of core promoter elements in B): INR, initiator; MTE, motif ten element; BREu, TFIIB recognition element upstream; DCE_S_I, downstream core element S-I; DCE_S_II, downstream core element S-II.

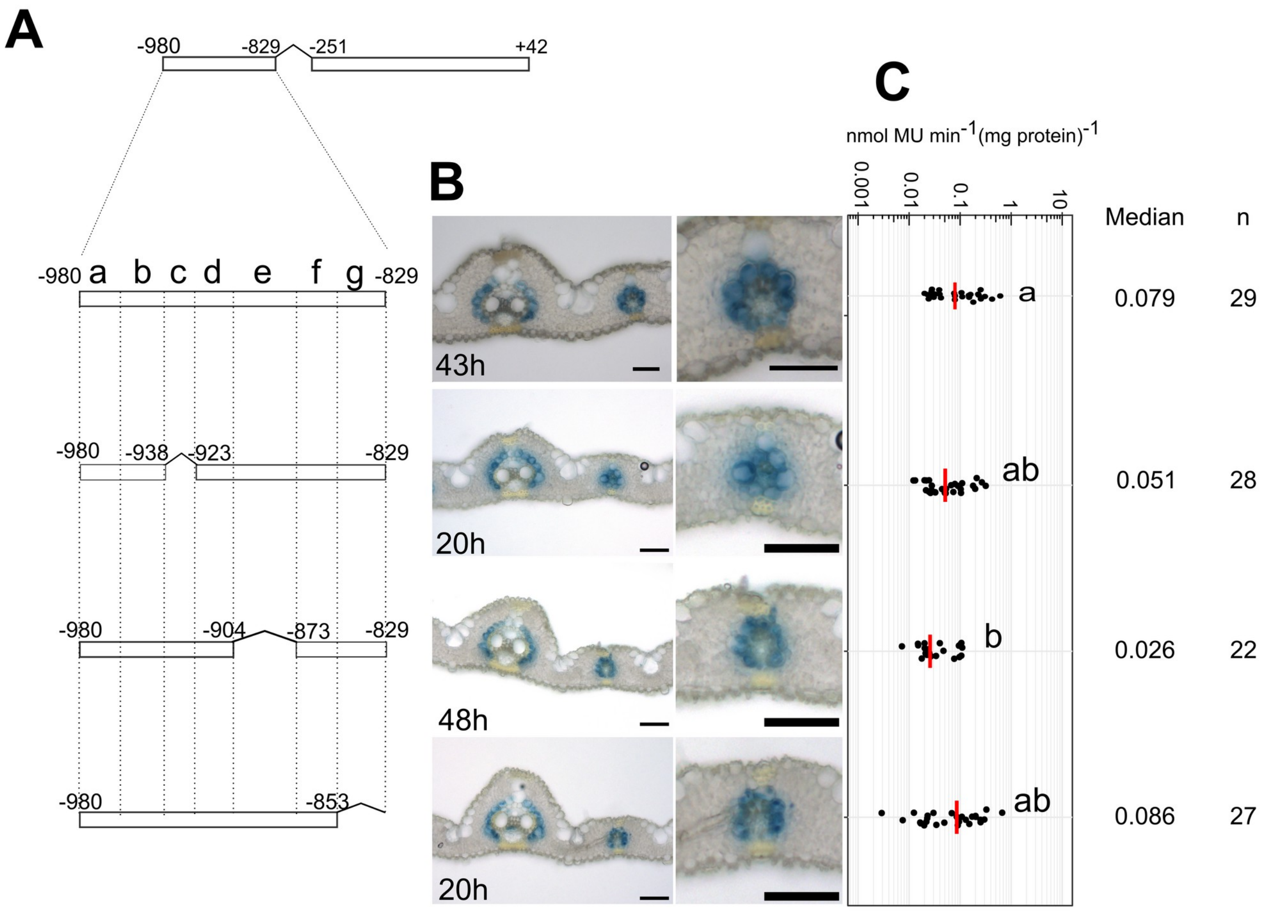


**Supplemental Figure 10. Subregions in distal enhancer not required for bundle sheath specific expression between nucleotides -980 to -829.** (**A**) Schematic of sequences fused to GUS reporter. (**B**) Representative images of leaf cross sections after GUS staining, zoomed-in images of lateral veins shown in right panels, scale bars = 50 um. (**C**) Promoter activity determined by the fluorometric 4-methylumbelliferyl-β-D-glucuronide (MUG) assay. Data subjected to pairwise Wilcoxon test with Benjamini-Hochberg correction. Lines with differences in activity that were statistically significant (adjusted *P*<0.05) labelled with different letters. Median catalytic rate of GUS indicated with red line, n indicates total number of transgenic lines assessed. n indicates total number of transgenic lines assessed.

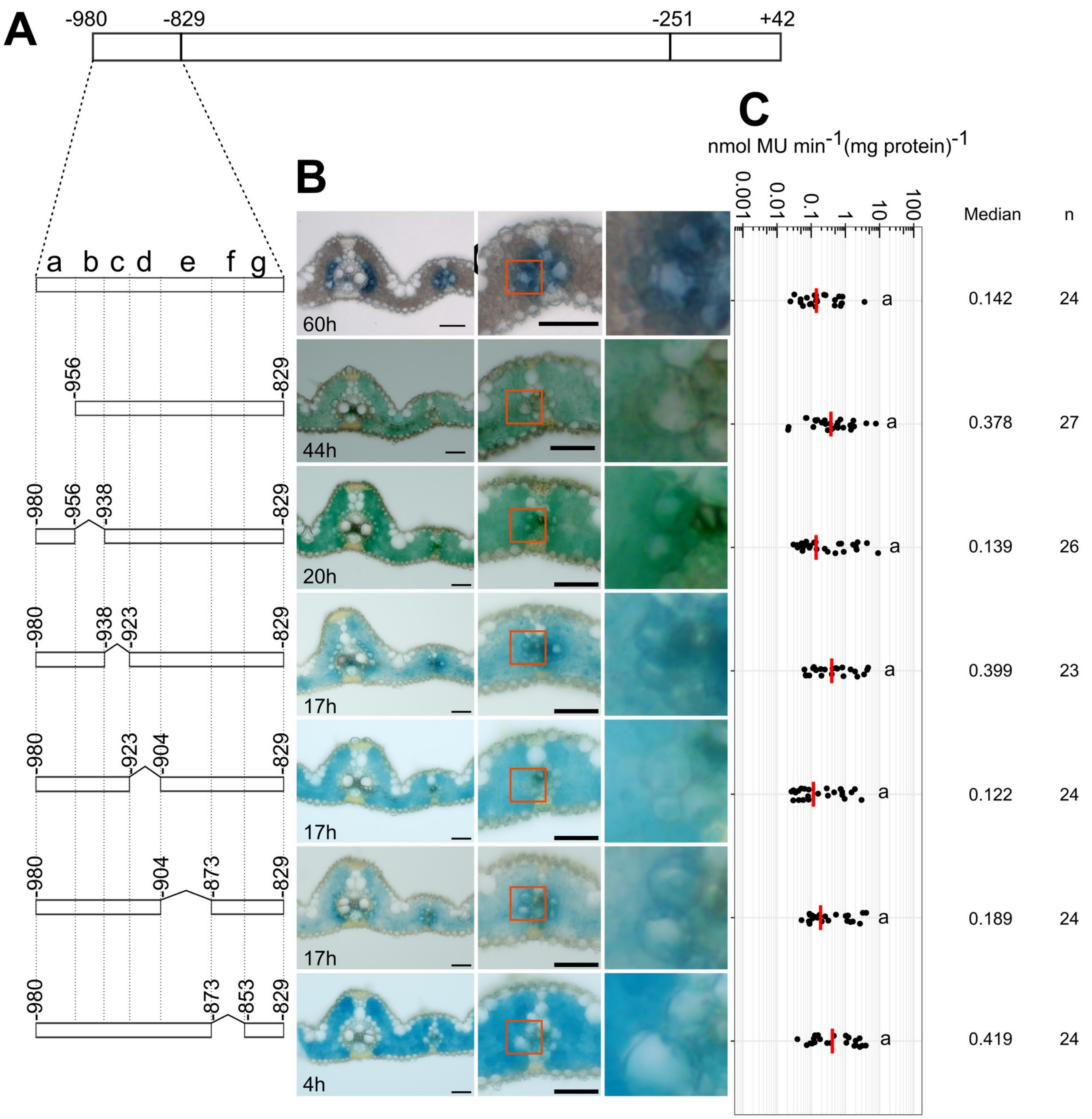


**Supplemental Figure 11. Regions between nucleotides -980 and -829 combined with nucleotides -828 to -252 repress mesophyll expression.** (**A**) Diagram showing deletions between nucleotides -980 and -829. (**B**) Representative images of leaf cross sections after GUS staining, zoomed-in images of lateral veins and bundle sheath cells shown in right panels, scale bars = 50 µm. (**C**) Promoter activity determined by the fluorometric 4-methylumbelliferyl-β-D-glucuronide (MUG) assay, Data subjected to pairwise Wilcoxon test with Benjamini-Hochberg correction. Lines with differences in activity that were statistically significant (adjusted *P*<0.05) labelled with different letters. Median catalytic rate of GUS indicated with red line, n indicates total number of transgenic lines assessed. n indicates total number of transgenic lines assessed.

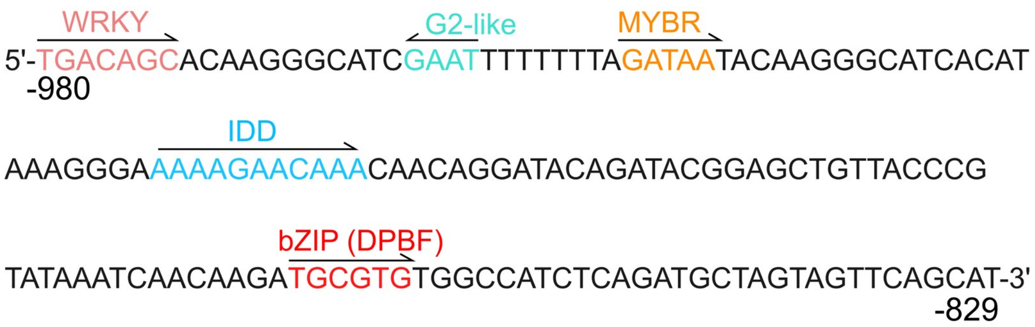


**Supplemental Figure 12. Nucleotide sequence of the distal enhancer. (A)** Transcription factor binding sites for WRKY, G2-like, MYBR, IDD, and bZIP highlighted, and orientation indicated.

| Subregion | Family | TF | MUS7 | motif |
|---|---|---|---|---|
| a | WRKY | WRKY1 | LOC_Os01g14440 | FIMO |
| a | WRKY | WRKY94 | LOC_Os12g40570 | FIMO |
| a | WRKY | WRKY121 | LOC_Os03g53050 | FIMO |
| a | G2like | GLK1 | LOC_Os06g24070 | FIMO |
| a | G2like | GLK2 | LOC_Os01g13740 | FIMO |
| a | G2like | MYS1 | LOC_Os12g01490 | FIMO |
| a | G2like | DLN168 | LOC_Os06g35140 | FIMO |
| a | G2like | MYR2 | LOC_Os03g03760 | FIMO |
| a | G2like | DLN52 | LOC_Os02g14490 | FIMO |
| a | G2like | HINGE1 | LOC_Os04g56990 | FIMO |
| a | G2like | DLN210 | LOC_Os08g33750 | FIMO |
| a | G2like | Os2R_MYB2 | LOC_Os01g04930 | FIMO |
| a | G2like | Os2R_MYB57 | LOC_Os05g37730 | FIMO |
| a | G2like | UCIP5 | LOC_Os12g39640 | FIMO |
| b | MYBR | MYBS1 | LOC_Os01g34060 | FIMO |
| b | MYBR | MYBS2 | LOC_Os10g41260 | FIMO |
| b | MYBR | MYBS3 | LOC_Os10g41200 | FIMO |
| b | MYBR | MYBR2 | LOC_Os08g04840 | FIMO |
| b | MYBR | MYBR3 | LOC_Os06g01670 | FIMO |
| b | MYBR | MYBR1 | LOC_Os01g09280 | FIMO |
| b | MYBR | MYBY2 | LOC_Os03g31230 | FIMO |
| b | MYBR | LHY | LOC_Os08g06110 | FIMO |
| d | IDD | IDD1 | LOC_Os03g10140 | FIMO |
| d | IDD | IDD2 | LOC_Os01g09850 | FIMO |
| d | IDD | IDD3 | LOC_Os09g38340 | FIMO |
| d | IDD | IDD4 | LOC_Os02g45054 | FIMO |
| d | IDD | IDD5 | LOC_Os07g39310 | FIMO |
| d | IDD | IDD6 | LOC_Os08g44050 | FIMO |
| d | IDD | IDD10 | LOC_Os04g47860 | FIMO |
| d | IDD | IDD11 | LOC_Os01g39110 | FIMO |
| f | SNAC | SNAC3 | LOC_Os01g09550 | FIMO |
| f | SNAC | NAC3 | LOC_Os07g12340 | FIMO |
| f | SNAC | NAC4 | LOC_Os01g60020 | FIMO |
| f | SNAC | NAC5 | LOC_Os11g08210 | FIMO |
| f | SNAC | NAC6 | LOC_Os01g66120 | FIMO |
| f | SNAC | NAC9 | LOC_Os03g60080 | FIMO |
| f | SNAC | NAP | LOC_Os03g21060 | FIMO |
| f | bZIP-A | bZIP9 | LOC_Os09g28310 | Kim et al., 2002 (A/CCACNNG,ACACNNG) |
| f | bZIP-A | bZIP10 | LOC_Os08g36790 | Kim et al., 2002 (A/CCACNNG,ACACNNG) |
| f | bZIP-A | bZIP11 | LOC_Os02g52780 | Kim et al., 2002 (A/CCACNNG,ACACNNG) |
| f | bZIP-A | bZIP3 | LOC_Os01g59760 | Kim et al., 2002 (A/CCACNNG,ACACNNG) |
| f | bZIP-A | bZIP4 | LOC_Os05g41070 | Kim et al., 2002 (A/CCACNNG,ACACNNG) |

**Supplemental Figure 13. Transcription factors used in transactivation effector assays.**

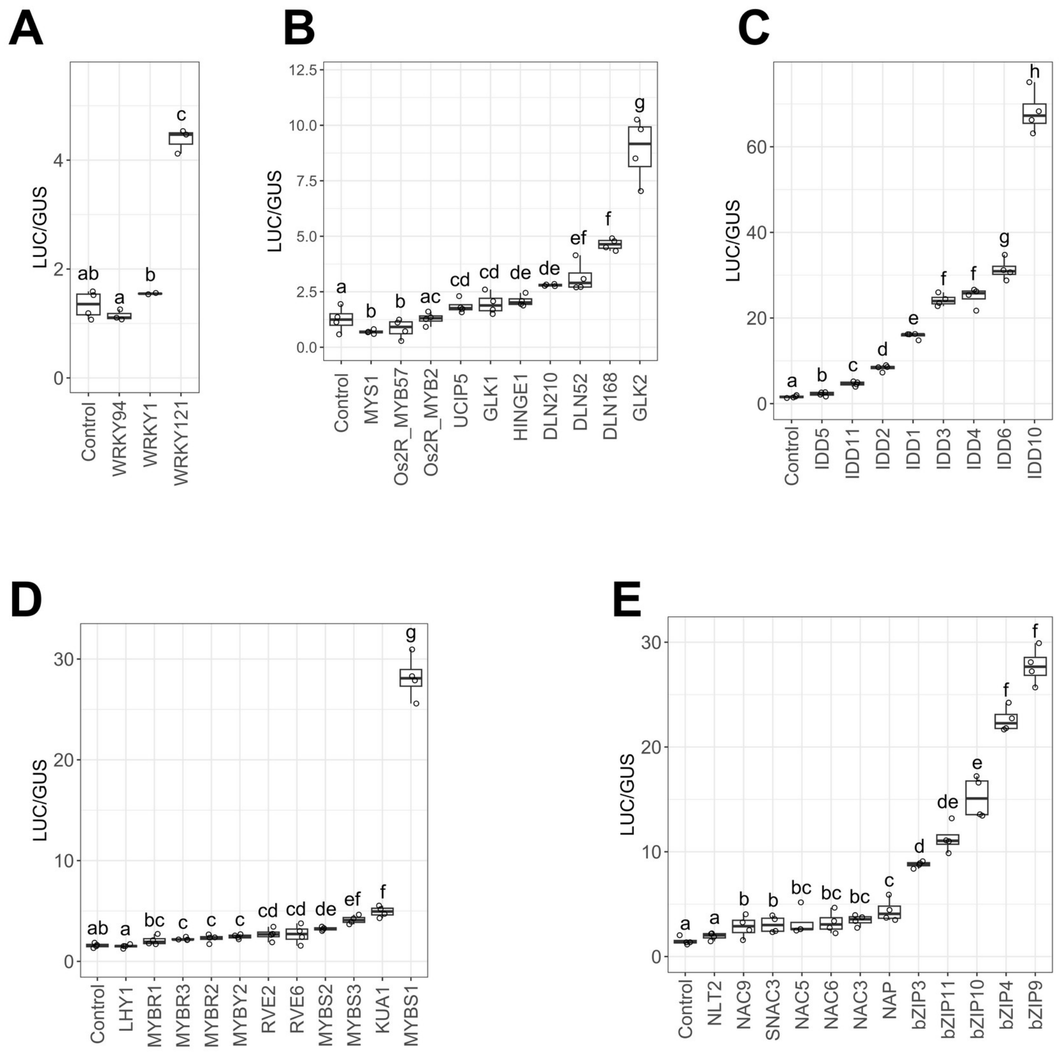


**Supplemental Figure 14. Effector assay showing the effect of WRKY (A), G2-like (B), IDD (C), MYB-related (D), SNAC and bZIP (E) transcription factors on outputs from the distal enhancer.** Data subjected to pairwise Wilcoxon test with Benjamini-Hochberg correction, samples with significantly different activities (adjusted *P*<0.05) are labelled with different letters.

**A**

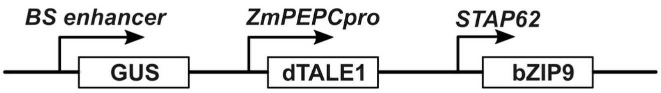
BS enhancer
ZmPEPCpro
STAP62
GUS
dTALE1
bZIP9

**B**

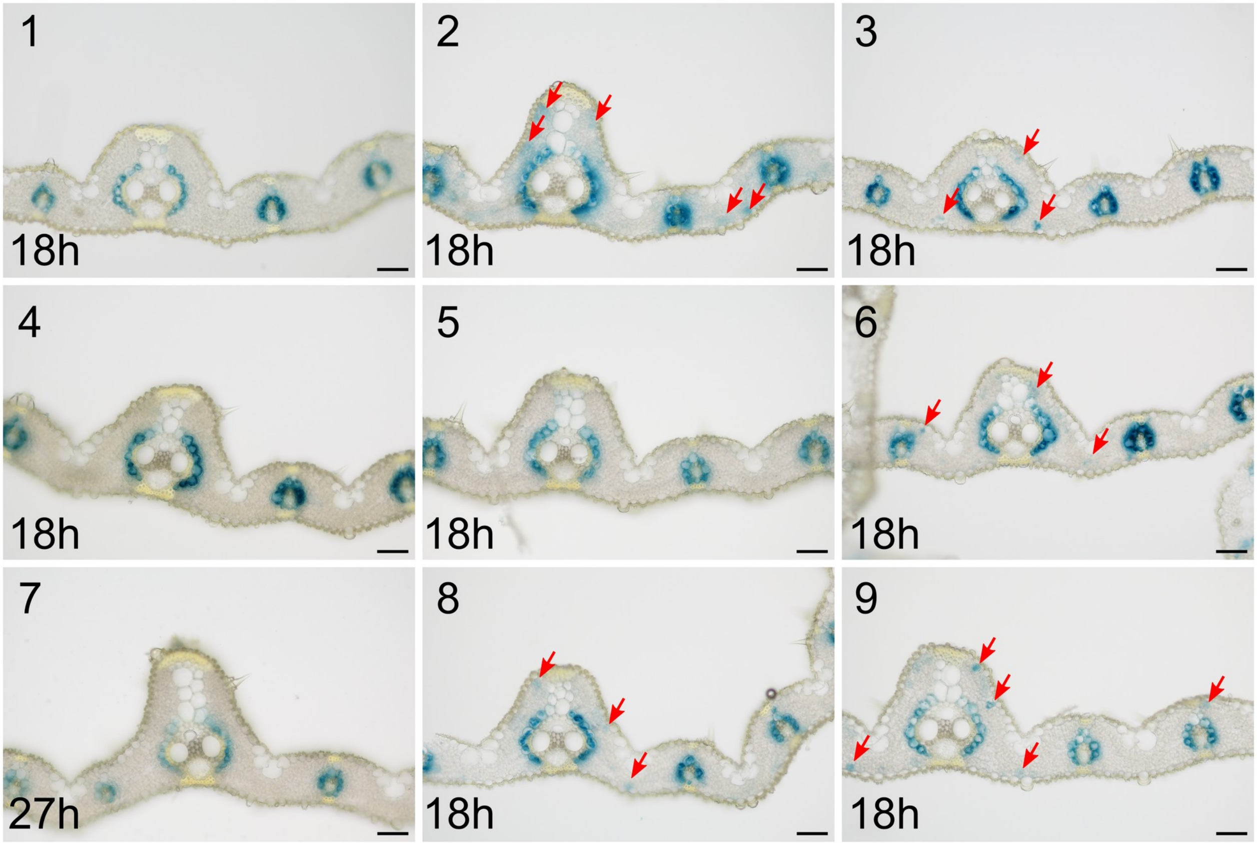
1
18h
2
18h
3
18h
4
18h
5
18h
6
18h
7
27h
8
18h
9
18h

C

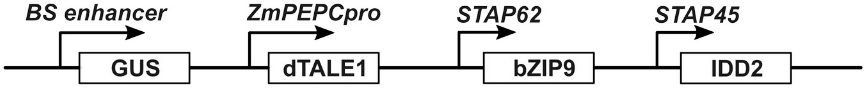

BS enhancer
ZmPEPCpro
STAP62
STAP45
GUS
dTALE1
bZIP9
IDD2


D

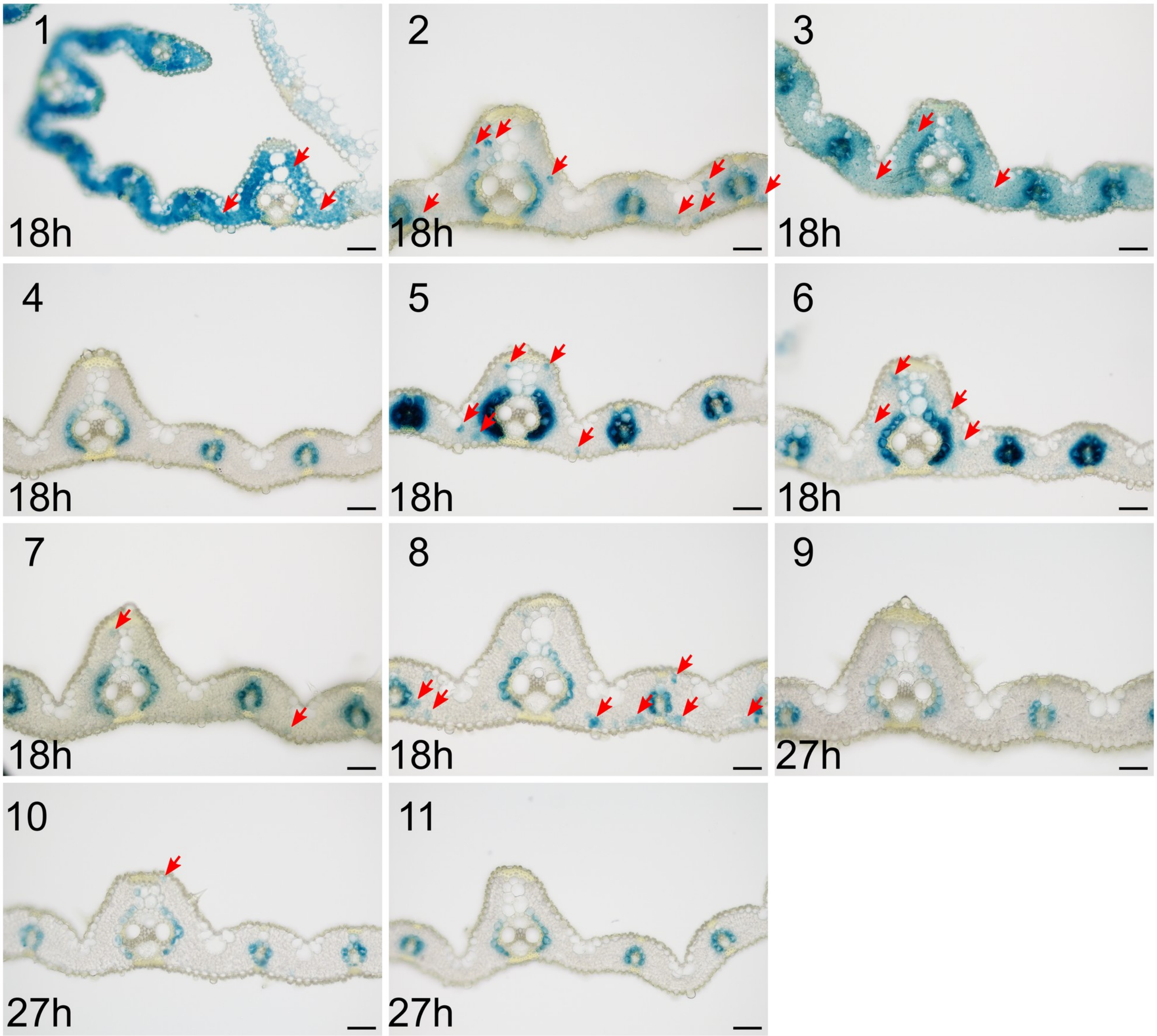

1
18h
2
18h
3
18h
4
18h
5
18h
6
18h
7
18h
8
18h
9
27h
10
27h
11
27h

E

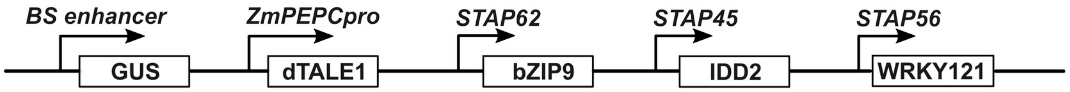


F

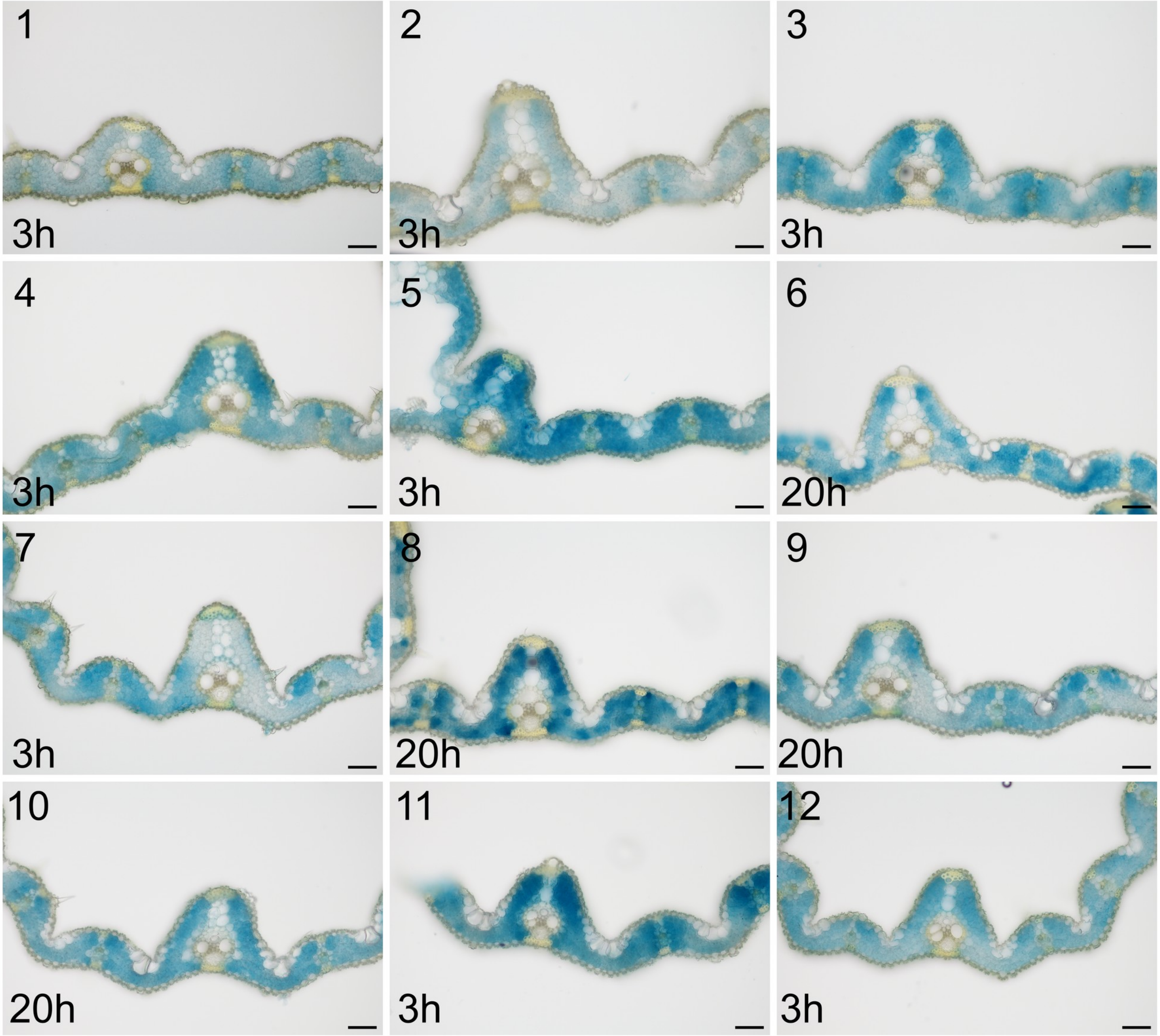


**Supplemental Figure 15. Impact of mis-expression of bZIP9, IDD2 and WRKY121 in mesophyll cells on GUS expression pattern driven by the bundle sheath enhancer. (A,C,E)** Schematics showing mis-expression of bZIP9, IDD2 and WRKY121 in mesophyll cells using the maize *PEPC* promoter driving dTALE1 in combination with STAP56, 45 and 62, the bundle sheath enhancer was fused with the *SiR* core promoter as a reporter. **(B,D,F)** Representative images of cross sections from transgenic lines after GUS staining, scale bars = 50 µm. Nine, eleven and twelve independent transgenic lines shown for each construct. Red arrows indicate GUS expressing mesophyll cells. Staining duration is displayed in the bottom-left corner, scale bars = 50 µm in B, D&F.

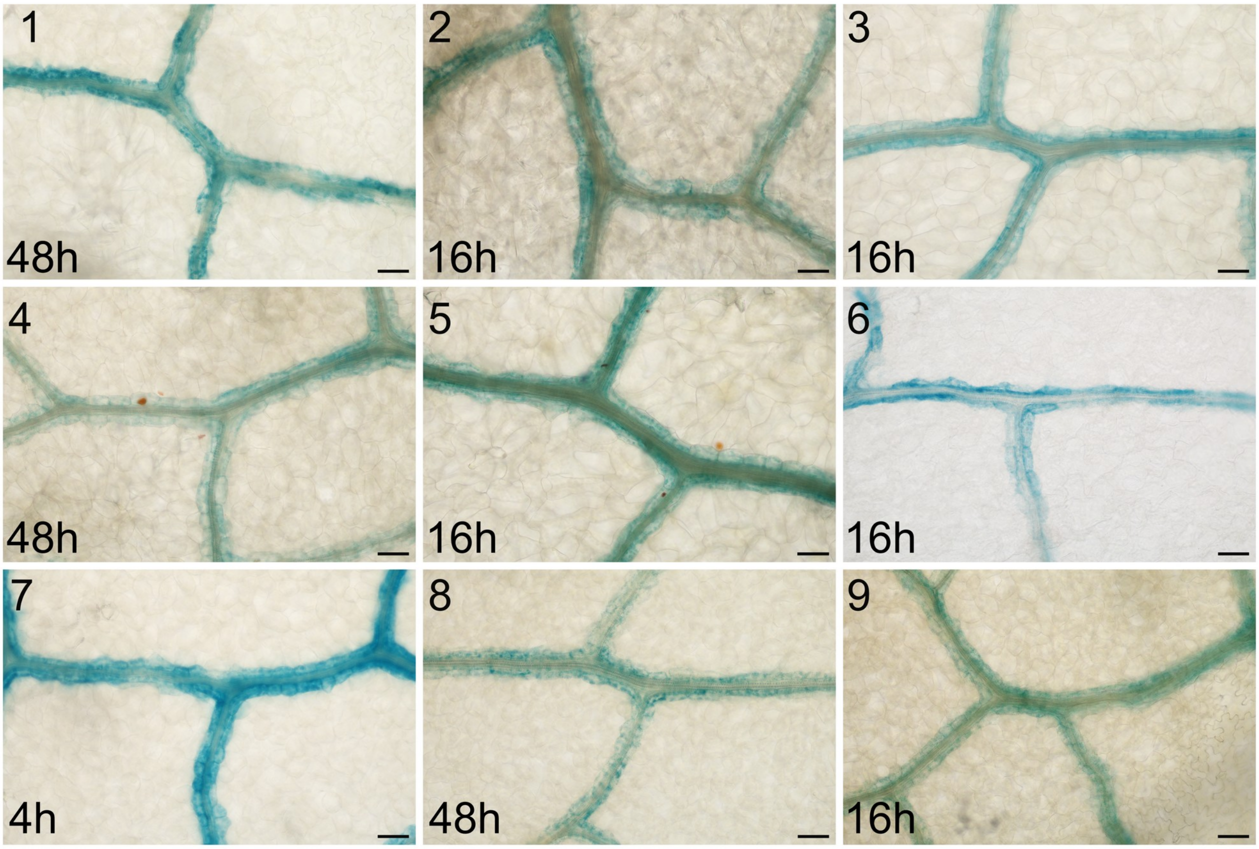


**Supplemental Figure 16. Three copies of the bundle sheath enhancer produce strong and specific bundle sheath specific expression in Arabidopsis.** Nine independent lines shown. Staining duration displayed in the bottom-left corner of each image, scale bars = 50 µm.

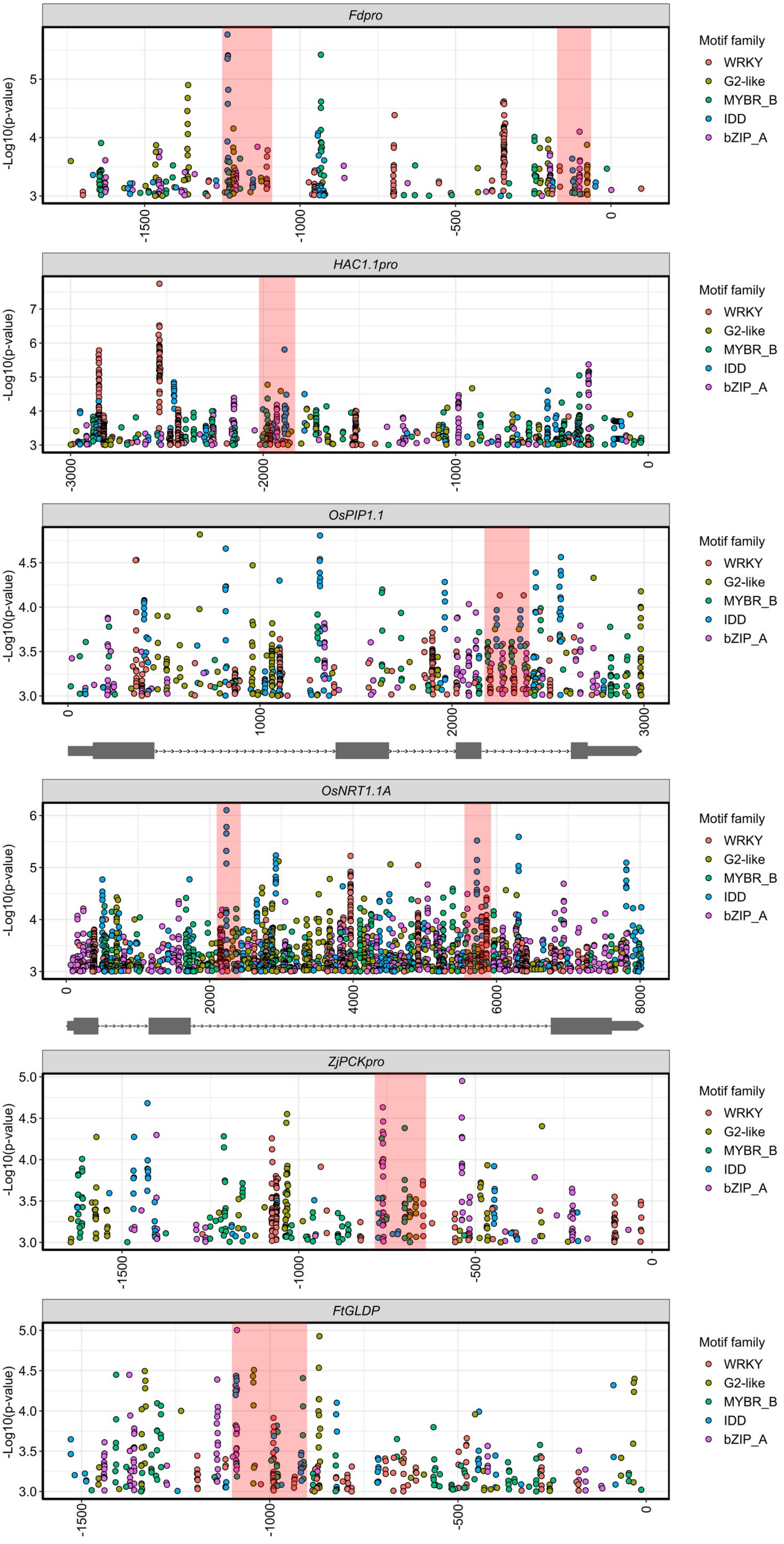


**Supplemental Figure 17. WRKY, G2-like, MYB-related, IDD and bZIP transcription factor binding sites identified by FIMO program in *Fd*, *HAC1;1*, *PIP1.1*, *NRT1.1A*, *ZjPCK* and *FtGLDP* promoters, regions <300 bp containing all five motif families highlighted in red.**
